# *Snhg26* long non-coding RNA regulates pluripotent cell states through a SINEB2-derived sequence

**DOI:** 10.64898/2026.05.21.726207

**Authors:** Victoire Fort, Gabriel Khelifi, Samer M.I. Hussein

## Abstract

Long-non coding RNAs (lncRNAs) are now well established players in gene expression regulation, but their detailed molecular mechanisms of action and underlying regulating sequences remain poorly understood. An emerging concept supports the idea that repeated sequences, more notably sequences derived from transposable elements (TEs), contribute to functional domains of lncRNAs. Here, we undertake the characterization of the function, interactors and functional domains of *Snhg26,* a lncRNA involved in reprogramming towards induced pluripotent stem cells (iPSCs) and maintenance of the pluripotent state of embryonic stem cells (ESCs). First, we show that modulation of *Snhg26* expression levels affects expression and splicing of genes involved in pluripotency and chromatin remodeling during the early steps of reprogramming and in ESCs. We also find that down-regulation of *Snhg26* increases the expression of SINEB2 transposable elements. Moreover, we identify hundreds of transcripts directly interacting with *Snhg26* with a significant enrichment of RNAs containing SINEB2 elements. Strikingly, loss of a SINEB2 sequence embedded within *Snhg26* abolishes its function in regulating pluripotent states. Our results thus support the idea that TEs constitute a source of functional units for lncRNAs and encourages further efforts to explore this concept.

## Introduction

Gene expression regulation is key to an organism’s development and homeostasis, and its disruption may cause several diseases, including developmental disorders and cancer^1^. Although pioneering studies of gene regulation have primarily focused on protein-coding genes, it is now recognized that the mammalian genome encodes thousands of long non-coding RNAs (lncRNAs), implicated in development and in disease progression as well^2–4^. High-throughput technologies have enabled better identification and annotation of those lncRNAs^5,6^, and functional characterization demonstrated that they affect gene regulation at multiple levels such as transcription, post-transcriptional processing, and translation^7,8^. They are also involved in subcellular localization of proteins and participate in the molecular structure of cellular organelles and nuclear subcompartments^7,8^. Such a diversity in lncRNA possible localizations and roles raised the need to discriminate common patterns within lncRNA primary sequence features or secondary structures that could serve as functional domains common to groups of lncRNAs^6^. This would allow the establishment of a lncRNA classification, which is so far based on their biogenesis (i.e. enhancer/promoter, intronic, sense/antisense, intergenic or bidirectional RNA), structure (i.e. linear or circular RNA), cellular localization or mode of action (i.e. in *cis* or *trans*)^7–12^. More importantly, identification of lncRNA domains would constitute a valuable tool to predict lncRNA functions, as is the case for protein-coding genes^13,14^. Furthermore, systematically establishing a list of lncRNA molecular interactors would also be helpful to group lncRNAs that have common interactomes and compare their functions. However, complete identification of interacting molecular partners is often lacking, and pinpointing features of the lncRNA sequence that are functionally relevant remains a challenge^8^.

In the last decade, a correlation between the presence and conservation of repeated sequences within lncRNA genes has led to the repeat insertion domains of lncRNAs (RIDL) hypothesis, according to which exonic transposable elements (TEs) provide roles for lncRNAs, thus constituting functional domains^15,16^. Transposable elements are sequences within genomic DNA that can replicate themselves independently of the host DNA and “jump” from a locus to another through copy-paste or cut-paste mechanisms. They participate to genome evolution and represent about half of the human genome. However, only a small fraction of TEs is still active and several TEs end up being transcribed as part of the gene they inserted into^17^. If about 0.3% of human mRNA possess a sequence derived from a TE, this number raises to 70-80% in lncRNAs^16,18–21^. The RIDL hypothesis thus postulated that lncRNA-embedded TEs could affect their regulation, localization and function. Examples include mouse SINEB2 and human free right *Alu* monomer repeat elements within antisense lncRNAs called SINEUPs. Those were shown to induce post-transcriptional expression up-regulation of their corresponding sense mRNAs^22,23^. Alternatively, presence of complementary SINE or *Alu* elements in a lncRNA and the 3’UTR of an mRNA can lead to intermolecular hybridization into a double-strand RNA complex triggering Staufen1-mediated mRNA decay^24,25^. These studies demonstrate the importance to study lncRNAs domains as they can unveil families of lncRNAs sharing similar sequence features with clear functional outcomes.

We previously identified a new nuclear lncRNA, *Snhg26*, highly expressed during embryonic development and in embryonic stem cells (ESCs) (Fort *et al.*, in preparation). *Snhg26* is responsible for increasing the number of induced pluripotent stem cells (iPSCs) during reprogramming of mouse somatic cells. In the same study, we also demonstrated its importance to maintain the pluripotent state of mouse and human ESCs. However, the underlying mechanism of action was not explored, and *Snhg26* interaction partners as well as its functional domains remain to be elucidated.

In the present study, we show that *Snhg26* lncRNA promotes pluripotency acquisition by up-regulating genes involved in mesenchymal-to-epithelial transition, pluripotency signaling pathways and chromatin remodeling during the early steps of reprogramming. Knock-out (KO) of *Snhg26* in mouse ESCs demonstrates that its expression is essential for *naïve* pluripotency maintenance. Interestingly, partial or total depletion of *Snhg26* in ESCs leads to modulations of isoform diversity, increases intron retention and affects splicing of pluripotency and chromatin remodeling-associated genes. This suggests that it is involved in their post-transcriptional regulation. We discovered that despite its nuclear localization, *Snhg26* does not bind the genome but rather binds to hundreds of RNAs, mostly involved in embryonic pattern specification. Inspection of *Snhg26* exonic sequence revealed presence of 4 TEs. When evaluating enrichment of *Snhg26* RNA partners for different TEs families, we observe a significant enrichment for SINEB2 sequences. Following *Snhg26* knock-down (KD) in ESCs, SINEB2 RNAs are found up-regulated transcriptome-wide, suggesting a role for *Snhg26* in post-transcriptional destabilization of these RNAs. Finally, overexpression during reprogramming to pluripotency of a *Snhg26* transcript lacking its SINEB2 sequence (*Snhg26*ΔSINEB2) abrogates enhancement of reprogramming. Moreover, no homozygous *Snhg26* KO-ESCs could be obtained when *Snhg26* is rescued by ectopic expression of a *Snhg26*ΔSINEB2 transgene. Together, our investigations reveal that the SINEB2 element within *Snhg26* sequence is essential to mediate its function in regulating the pluripotent state, contributing to the validation of the RIDL hypothesis.

## Results

### 1. *Snhg26* lncRNA overexpression accelerates acquisition of pluripotency hallmarks early during reprogramming while loss of *Snhg26* promotes exit of the *naïve* pluripotency state

In order to characterize the effect of *Snhg26* on the reprogramming process, we performed long-read RNA sequencing (lrRNAseq) whole transcriptome analysis during the reprogramming of mouse neural progenitor cells (NPCs) into iPSCs, as we previously described in Fort *et al.* (in preparation). We compared the consequences of overexpressing the *Oct4, Klf4* and *Sox2* (OKS) reprogramming factors along with either a doxycycline (DOX)-inducible transgene carrying the cDNA of *Snhg26* lncRNA or an inert RNA of similar length transcript (i.e. the antisense to the *Luciferase* gene, “control”) at different timepoints of the process (D2, D4, D6, D10, D14, D20 in FBS-LIF + DOX) (**Fig. 1A** and **Table S1**). NPC lines (D0) as well as cells at D20 in 2i-LIF (corresponding to the endpoint iPSC stage) were used as controls. To get a global view of the routes taken by the reprogramming cell lines, we performed multidimensional scaling analysis of global gene expression (**Fig. 1B**). Even though trajectories of OKS+control and OKS+*Snhg26* were globally similar, when focusing on the first half of the process (from D2 to D14), a major difference was observable, with trajectories splitting their ways before rejoining by the end of reprogramming (D20). This suggested that *Snhg26* lncRNA overexpression affects global gene expression during the first half of reprogramming towards pluripotency, which is consistent with our previous findings of *Snhg26* increasing the number of cells engaging into reprogramming (Fort *et al.*, in preparation). Next, we compared the expression profiles of specific subsets of genes that act as hallmarks of reprogramming steps^26^ (Fort *et al.*, in preparation) (**Table S1**). We found that OKS+*Snhg26* reprogramming showed some dissimilarly to OKS+control for some gene sets (**Fig. 1C** and **Extended Data Fig. 1A**). *Snhg26* overexpression during reprogramming induced earlier and higher activation of immediate, intermediate, early ESC-like and ESC-like genes, compared to the control condition. More particularly, MET genes (genes associated with mesenchymal-to-epithelial transition) were activated from the beginning of reprogramming, while this happened only from D4 in OKS+control (**Fig. 1C**). This was accompanied by decreased expression of epithelial-to-mesenchymal genes starting at D4 in OKS+*Snhg26* reprogramming and only from D10 in control condition (**Extended Data Fig. 1A**), suggesting that *Snhg26* accelerates the MET transition, which is an essential step of reprogramming to iPSCs^27,28^.

**Figure 1:**
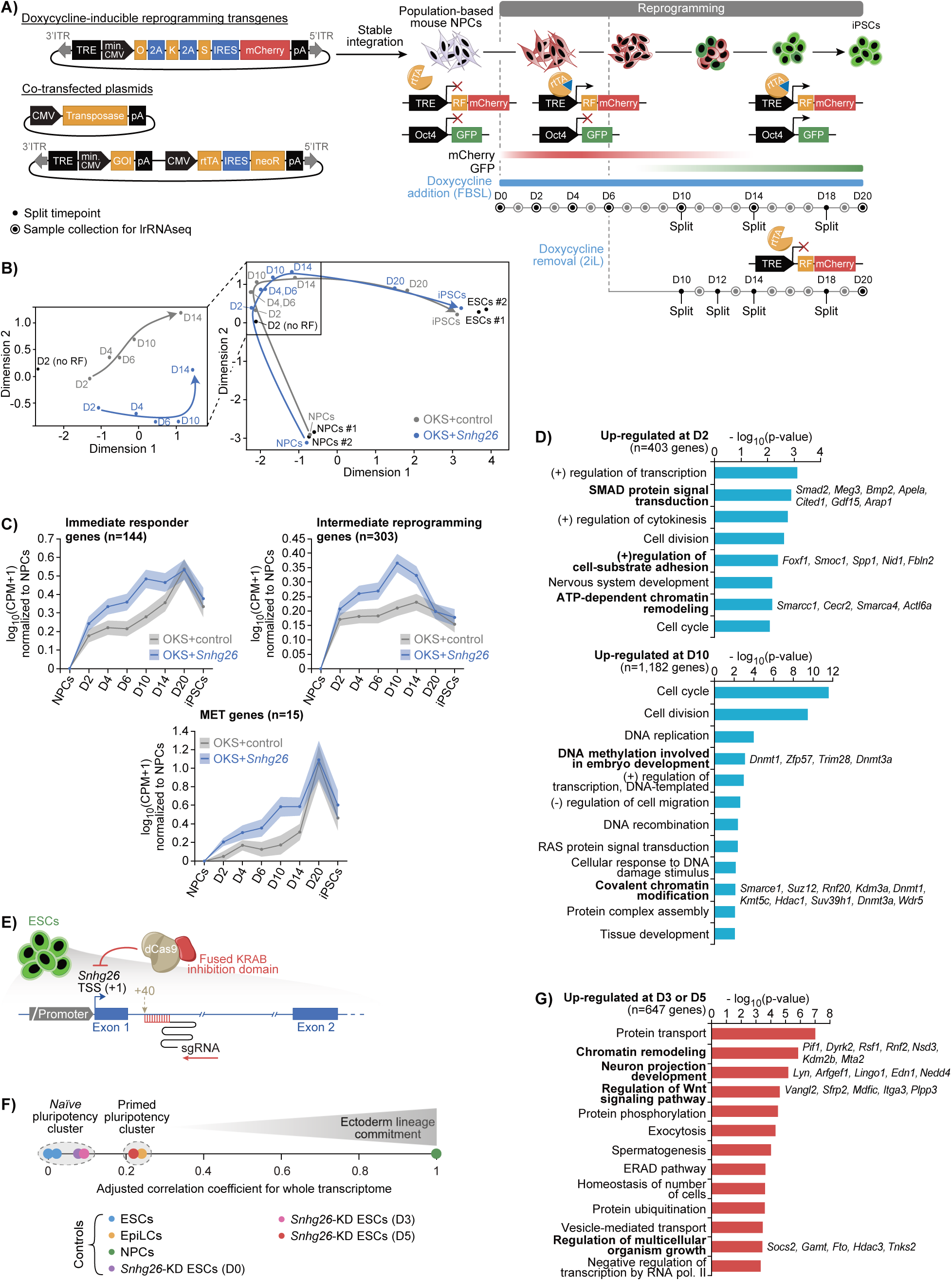
*Snhg26* lncRNA overexpression accelerates acquisition of pluripotency hallmarks early during reprogramming while loss of *Snhg26* promotes exit of the *naïve* pluripotency state. **A)** Schematic overview of the experimental reprogramming pipeline. For more details refer to Fort *et al.* (in preparation). ITR: Inverted Terminal Repeat; TRE: Tet Response Element; min. CMV: minimal CMV promoter; IRES: Internal Ribosome Entry Site; pA: poly(A) signal sequence; GOI: Gene Of Interest, i.e. *Snhg26* or the antisense to the *Luciferase* gene (control); rtTA: reverse tetracycline-controlled TransActivator; neoR: neomycin Resistance gene; lrRNAseq: long-read RNA sequencing; NPCs: Neural Progenitor Cells; iPSCs: induced Pluripotent Stem Cells; RF: Reprogramming Factors (i.e. *Oct4* (O), *Klf4* (K), *c-Myc* (M) and *Sox2* (S)); FBSL: FBS-LIF medium; 2iL: 2 inhibitors + LIF medium. **B)** Multidimensional scaling plot of the distances between OKS+control (grey) and OKS+*Snhg26* (blue) reprogramming samples between D0 (NPC samples) and final iPSCs, with ESCs as controls (right). A close-up of samples between D2 and D14 is also shown (left). ESCs: Embryonic Stem Cells. **C)** Plots depicting the logged fold change in expression in read counts as counts per million (CPM) of immediate responder (n=144) and intermediate reprogramming (n=303) genes (gene lists from Hussein *et al.*, Nature 2014), or mesenchymal-to-epithelial transition (MET) (n=15) genes in each timepoint of the OKS+control (grey) or OKS+*Snhg26* (blue) reprogramming, normalized to NPCs. Lines represent the average expression value at each timepoint and the shaded area represents the standard error of the mean (SEM). The data for OKS+control is the same as in Fort *et al.* (in preparation). Also see **Extended Data Fig. 1A**. **D)** Bar graphs depicting the logged p-value of GO-terms of enriched biological processes for which genes undergo up-regulation at D2 (n=403 genes, top) and D10 (n=1,182 genes, bottom) of OKS+*Snhg26* reprogramming. P-value < 0.01. (+)/(-): Positive/Negative. Also see **Extended Data Fig. 1B**. **E)** Schematic of the CRISPR-interference system used to knock-down expression of *Snhg26* lncRNA in mouse ESCs. dCas9: dead Cas9; TSS: Transcription Start Site; sgRNA: single-guide RNA. **F)** Representation of the adjusted correlation coefficient between the gene expression datasets of ESCs with knock-down (KD) of *Snhg26* lncRNA for 0 (n=1), 3 (n=1) or 5 (n=1) days and wild-type ESCs (n=2), EpiLCs (n=1) or NPCs (n=2). EpiLCs: Epiblast-Like Cells. **G)** Bar graph depicting the logged p-value of GO-terms of enriched biological processes for which genes undergo up-regulation at D3 and D5 (n=647 genes) of *Snhg26* KD. FDR ≤ 0.1. pol.: polymerase.

To investigate differences between the two reprogramming conditions, we first analyzed which biological processes were enriched among the differentially expressed genes (DEGs) (**Table S1**). We focused on DEGs between D2 and D14 as these were the timepoints were most of the changes occur, as seen previously (**Fig. 1B**). GO-terms enrichment analysis showed that the genes up-regulated at D2 and D10 of OKS+*Snhg26* compared to OKS+control reprogramming were associated to processes such as “SMAD protein signal transduction” (e.g. *Smad2, Bmp2, Gdf15*), which is linked with the regulation of pluripotency genes expression^29^ and embryonic development^30,31^ (**Fig. 1D** and **Table S1**). Several GO-terms corresponded to regulation of cell division (e.g. *Map10, Aurkb, Bub1b*) and cell-substrate adhesion (e.g. *Fbln7, Fblim1, Hepacam*), which was coherent with MET genes being up-regulated earlier when *Snhg26* is overexpressed (**Fig. 1D**, **Extended Data Fig. 1B-C** and **Table S1**). Genes involved in “cellular response to TGF-ß stimulus” and “negative regulation of BMP signaling pathway” were down-regulated early on during reprogramming of OKS+*Snhg26*. Moreover, activators of the MAPK transduction pathway were down-regulated at D14 (**Extended Data Fig. 1C** and **Table S1**). Regulation of these molecular signaling pathways was shown to favor populations of mouse ESCs with high *Nanog* expression levels and to promote self-renewal of pluripotent cells^32^. Biological processes indicative of loss of differentiated cell identities (e.g. “multicellular organism development”, “cell fate commitment”) (**Extended Data Fig. 1C** and **Table S1**) were also enriched in down-regulated genes, further demonstrating the conversion of the NPCs into iPSCs. Altogether, these results suggested that *Snhg26* expression accelerates the dynamics of the reprogramming process and enhances the commitment of cells towards pluripotency establishment, consistent with previous published observations (Fort *et al.*, in preparation). Surprisingly, genes involved in DNA methylation (e.g. *Dnmt1* and *Dnmt3a*) and ATP-dependent chromatin remodeling (e.g. *Smarcc1, Cecr2, Smarca4*) were also enriched at D2 and D10 (**Fig. 1D** and **Table S1**). In OKS+control, average expression of those genes decreased to then increase at D4 while in OKS+*Snhg26* they increased as early as D2 (**Extended Data Fig. 1D** and **Table S1**). Most of these were positive regulators of the chromatin. For instance, *Kdm6a* is an important regulator of reprogramming, known to open up the chromatin by demethylating H3K27me3 marks^33,34^. *Kdm3a* demethylates H3K9 to promote transcription activation, *Smarcc1* and *Smarca4* are also transcriptional activators. Enrichment in these types of genes may thus lead to the acceleration of the reprogramming kinetics observed when overexpressing *Snhg26*, as chromatin marks are well-known gatekeepers of the cell state^35–37^. Finally, we observed a significant up-regulation as early as D2 of imprinted genes *Rian* and *Meg3*, which were demonstrated as pivotal to obtain *bona fide* iPSCs^38,39^ (**Extended Data Fig. 1D** and **Table S1**). Altogether, these results suggest that *Snhg26* enhances reprogramming through up-regulation of genes involved in pluripotency signaling pathways and chromatin remodeling, allowing earlier commitment of cells towards reestablishment of the pluripotent state.

On top of its role in accelerating reprogramming to pluripotency, we also previously showed that down-regulating *Snhg26* expression levels in mouse and human ESCs led to a decrease of pluripotency genes (Fort *et al.*, in preparation). To better understand the effect of *Snhg26* KD on the pluripotency-regulatory network, we submitted samples of CRISPR interference-mediated *Snhg26* KD to lrRNAseq (**Fig. 1E** and **Table S1**). We used an adjusted Pearson correlation to compare gene expression patterns of *Snhg26-*KD ESCs after 0, 3 and 5 days of KD to those of mouse ESCs (i.e. pluripotent stem cells with a *naïve* state), Epiblast-Like Cells (EpiLCs) (i.e. pluripotent stem cells more primed to differentiate than ESCs) and NPCs (progenitor cells from the ectodermal lineage) (**Fig. 1F**). With this, we found that global gene expression from D0 and D3 *Snhg26-*KD cells correlated better with ESCs. However, D5 *Snhg26-*KD cells gene expression was closer to expression patterns found in EpiLCs. This result suggested that despite retaining their pluripotent identity, knock-down of *Snhg26* lncRNA expression in ESCs primed those cells for differentiation.

Next, we performed GO-terms enrichment analysis on the genes that were significantly up- or down-regulated in the D3 and D5 *Snhg26-*KD ESCs (n=1,169) compared to control samples (**Fig. 1G**). No enrichment was found for genes significantly down-regulated. However, up-regulated genes (n=647) were associated with chromatin remodeling and biological processes related to differentiation processes, such as “neuron projection development” and “regulation of multicellular organism growth” (**Fig. 1G** and **Table S1**). Regulation of the canonical Wnt signaling pathway also indicated a transition from a *naïve* to a primed pluripotent state^40^. Altogether, these results suggested ESC differentiation, further validating our Pearson analysis and previous observations that *Snhg26* is required to maintain *naïve* pluripotency (Fort *et al.*, in preparation).

### 2. *Snhg26* lncRNA expression is essential for *naïve* pluripotency maintenance

Because genes and signaling pathways implicated in embryonic development and differentiation were enriched in both *Snhg26* reprogramming and *Snhg26*-KD lrRNAseq analyses, and because we showed that *Snhg26* was required for *naïve* pluripotency maintenance, we wondered whether it would be essential to support pluripotency of mouse ESCs. To do so, we decided to knock-out the last exon of *Snhg26*, which represents 92.5% of its whole exonic sequence using CRISPR-Cas9 (**Fig. 2A** and **Table S2**). However, because KD experiments showed that *Snhg26* is required and may be necessary to maintain the pluripotent state of ESCs, we performed the KO while concomitantly and stably integrating a DOX-inducible construct allowing expression of full-length *Snhg26*. An antisense of the *Luciferase* gene was used as a control. We derived ESC KO clones and quantified the number of clones obtained from KO+control and KO+*Snhg26* ESCs using standard PCR-based amplification of *Snhg26* gene locus (**Extended Data Fig. 2A-B** and **Table S2**) and assessment of endogenous and transgenic *Snhg26* copy number by qPCR (**Extended Data Fig. 2C-D** and **Table S2**). Consistent with the role of *Snhg26* in sustaining pluripotency, we were able to achieve complete removal of both alleles (-/-) and heterozygous deletion (+/-) of exon 4 with a higher frequency when the full-length *Snhg26* lncRNA was overexpressed, compared to the control condition (**Fig. 2B**). This was validated in several ESC lines from different genetic backgrounds (C57BL/6 X 129/sv for R1 and Oct4-GFP (O4G) lines; C57BL/6 X 129/sv for V6.5 line), suggesting that our findings were not mouse background dependent (**Extended Data Fig. 2D** and **Table S2**).

**Figure 2:**
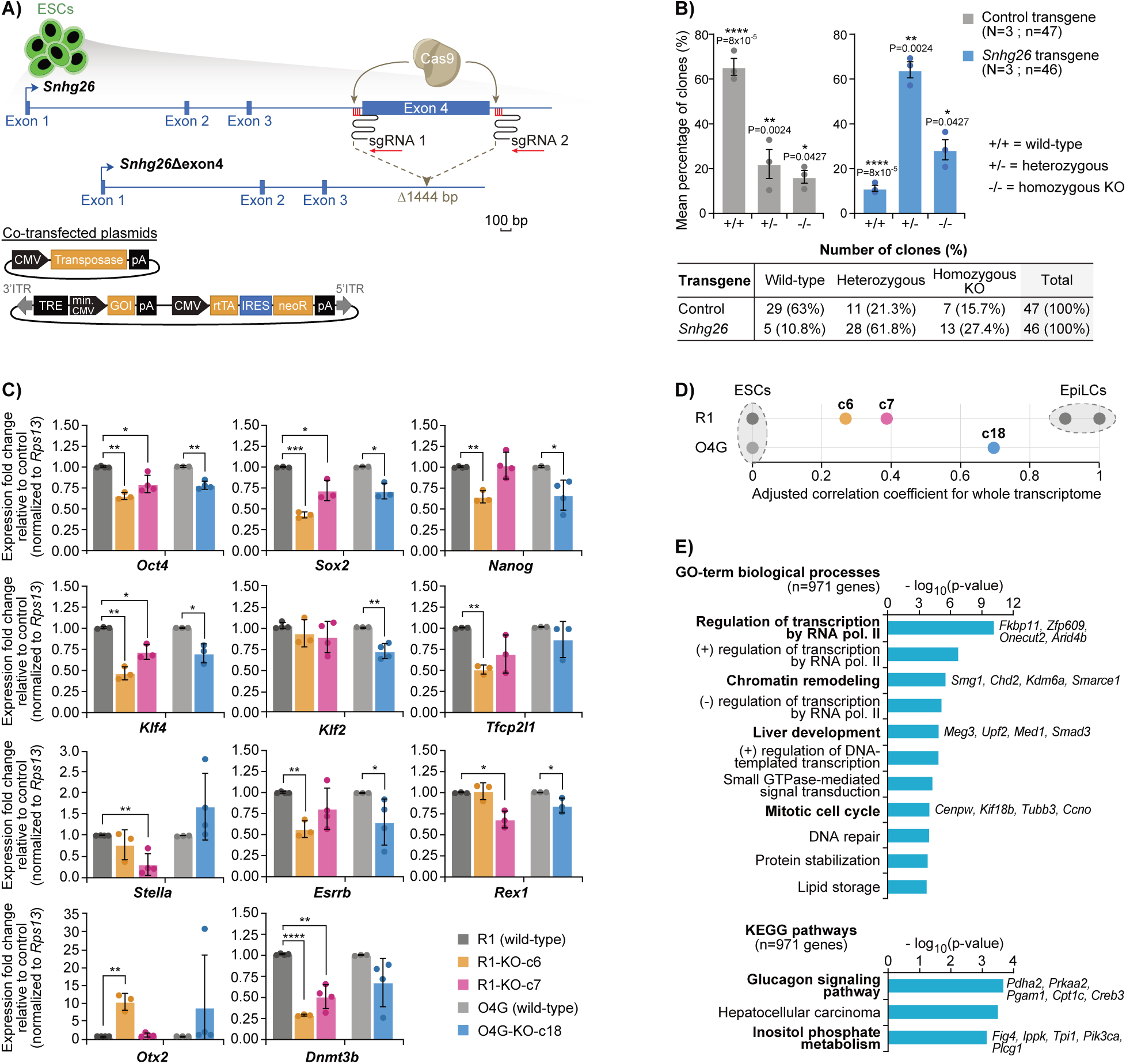
*Snhg26* lncRNA expression is essential for *naïve* pluripotency maintenance. **A)** Schematic of the CRISPR-Cas9 system used to knock-out the last exon of *Snhg26* lncRNA in mouse ESCs with stable integration of transgenes allowing inducible expression of genes of interest (GOI). bp: base pair. **B)** Bar graphs (top) showing the mean percentage of ESC clones obtained with a wild-type (+/+), heterozygous (+/-) or homozygous KO (-/-) locus of *Snhg26* gene, for the conditions expressing a control (grey; n=47) or *Snhg26* (blue; n=46) transgene. Error bars represent the SEM and each dot represents the replicates (N=3 ESC lines). P-values were calculated by one-tailed t-test. Table (bottom) indicates the number of clones (and the percentage they represent) obtained for each experiment. * < 0.05, ** < 0.01, **** < 0.0001. KO: Knock-Out. Also see **Extended Data Fig. 2D**. **C)** Bar graphs representing the mean expression fold change of *naïve* and formative pluripotency genes in KO-ESC clones relative to their corresponding wild-type ESC lines (R1 and O4G), normalized to *Rps13* housekeeping gene. Error bars represent the Standard Deviation (STD). P-values were calculated by one-tailed t-test with unequal variance. * < 0.05, ** < 0.01, *** < 0.001, **** < 0.0001. Also see **Extended Data Fig. 2E**. **D)** Representation of the adjusted correlation coefficient between the gene expression datasets of R1-KO-c6 (n=1), R1-KO-c7 (n=1), O4G-KO-c18 (n=1) ESC clones and wild-type R1 ESCs (n=1), wild-type O4G ESCs (n=1) and EpiLCs (n=2). **E)** Bar graphs depicting the logged p-value of GO-terms of enriched biological processes (top) or of KEGG pathways (bottom) for which genes undergo up- and down-regulation (n=971 genes) in KO-ESC clones compared to wild-type ESCs. FDR ≤ 0.1.

Core pluripotency genes *Oct4, Sox2* and *Nanog* expression decreased significantly in R1 and O4G KO-ESC clones (R1 clones c6 and c7; O4G clone c18, these clones do not contain the *Snhg26* transgene) (**Fig. 2C** and **Extended Data Fig. 2E**). Interestingly, *naïve* pluripotency markers *Klf4, Klf2, Tfcp2l1, Stella, Esrrb* and *Rex1,* as well as formative pluripotency markers *Otx2* and *Dnmt3b* also decreased in expression, except *Otx2* was significantly up-regulated in clone c6 (**Fig. 2C**). Next, to get a more comprehensive view of the effect of *Snhg26* KO on gene expression and pluripotency, we performed lrRNAseq on wild-type (WT) ESCs, EpiLCs and the KO-ESC clones (**Table S2**). By comparing the whole transcriptome of these different lines, we found that KO-ESC clones clustered in between ESCs and EpiLCs (**Fig. 2D**). GO-terms enrichment analysis on genes up- and down-regulated in KO-ESC clones (n=971 genes) compared to WT ESCs showed similarities with *Snhg26-*KD ESCs, with genes involved in “chromatin remodeling” and “liver development” being enriched (**Fig. 2E** and **Table S2**). Interestingly, KEGG pathway enrichment involved terms such as “glucagon signaling pathway” and “inositol phosphate metabolism”, suggesting a shift in the metabolism of KO-ESCs (**Fig. 2E** and **Table S2**). Taken together, these results suggested that KO-ESCs lose their *naïve* pluripotency state to resemble cells more advanced on the differentiation spectrum, and that *Snhg26* lncRNA is required to maintain the *naïve* pluripotent state, confirming the results from the KD experiments (**Fig. 1F-G**). These results also highlight more deeply the impact of *Snhg26* loss of expression on the pluripotency regulatory network, and reveal that *Snhg26* lncRNA is essential to sustain *naïve* pluripotency.

### 3. Loss of *Snhg26* impedes proper splicing of pluripotency and chromatin remodeling - associated transcripts

Taking advantage of our lrRNAseq datasets on both *Snhg26*-KD and KO-ESCs, we next assessed alternative splicing (AS), and isoform classification and diversity. Depletion (KD) of *Snhg26* lncRNA revealed an increase in transcripts that matched reference annotations (“Full Splice Matches” (FSM) or partially “Incomplete Splice Matches” (ISM) - 70.2% in *Snhg26-*KD ESCs vs 63.2% in control ESCs), along with a decrease in novel isoforms (from known splice junctions “Not-In-Catalogue” (NIC) and novel splice junctions “Novel-Not-in-Catalogue” (NNC) - 27.4% in *Snhg26-*KD ESCs vs 34.3% in control ESCs) (**Fig. 3A** and **Table S3**). Globally, there was 1.6 times more abundant isoforms in *Snhg26-*KD ESCs (n=570) than their control counterparts (n=359). An opposite observation was made for KO-ESCs, where there was less abundance in transcripts in KO-ESCs (n=381) than in WT ESCs (n=614). Accordingly, annotated isoforms declined (55.1% in KO-ESCs vs 58.2% in wild-type ESCs) while proportions of novel isoforms rose (41.7% in KO-ESCs vs 37.1% in wild-type ESCs) (**Fig. 3B** and **Table S3**). Irrespective of depletion (KD) or deletion (KO) strategies, these results nonetheless suggest that alterations in *Snhg26* levels in ESCs affects alternative splicing and the resulting isoform diversity. To further address this, we quantified the different types of AS events detected in ESCs with *Snhg26* depletion (n=56,033 events) or deletion (n=51,345 events). In *Snhg26-*KD ESCs, we found more AS events (n=1,308) than in control ESCs (n=801), with the majority of AS events corresponding to Alternative First exons (**Fig. 3C** and **Table S3**). This was in keeping with a 1.5 folds increase in the proportion of FSM isoforms with alternative 5’ ends in *Snhg26-*KD ESCs (**Fig. 3D** and **Table S3**). In contrast, there were less AS events in KO-ESCs (n=669) than in WT ESCs (n=952) (**Fig. 3C** and **Table S3**), in keeping with the previous result of isoform diversity. The only exception was for intron retention events that were more included in both KD- and KO-ESCs than in WT cells, suggesting a common phenomenon of intron retention. Together, these results show that *Snhg26* deletion or depletion leads to differential alternative splicing and favors isoform diversity, suggesting that *Snhg26* may be involved in regulation of alternative splicing and more specifically, intron retention.

**Figure 3:**
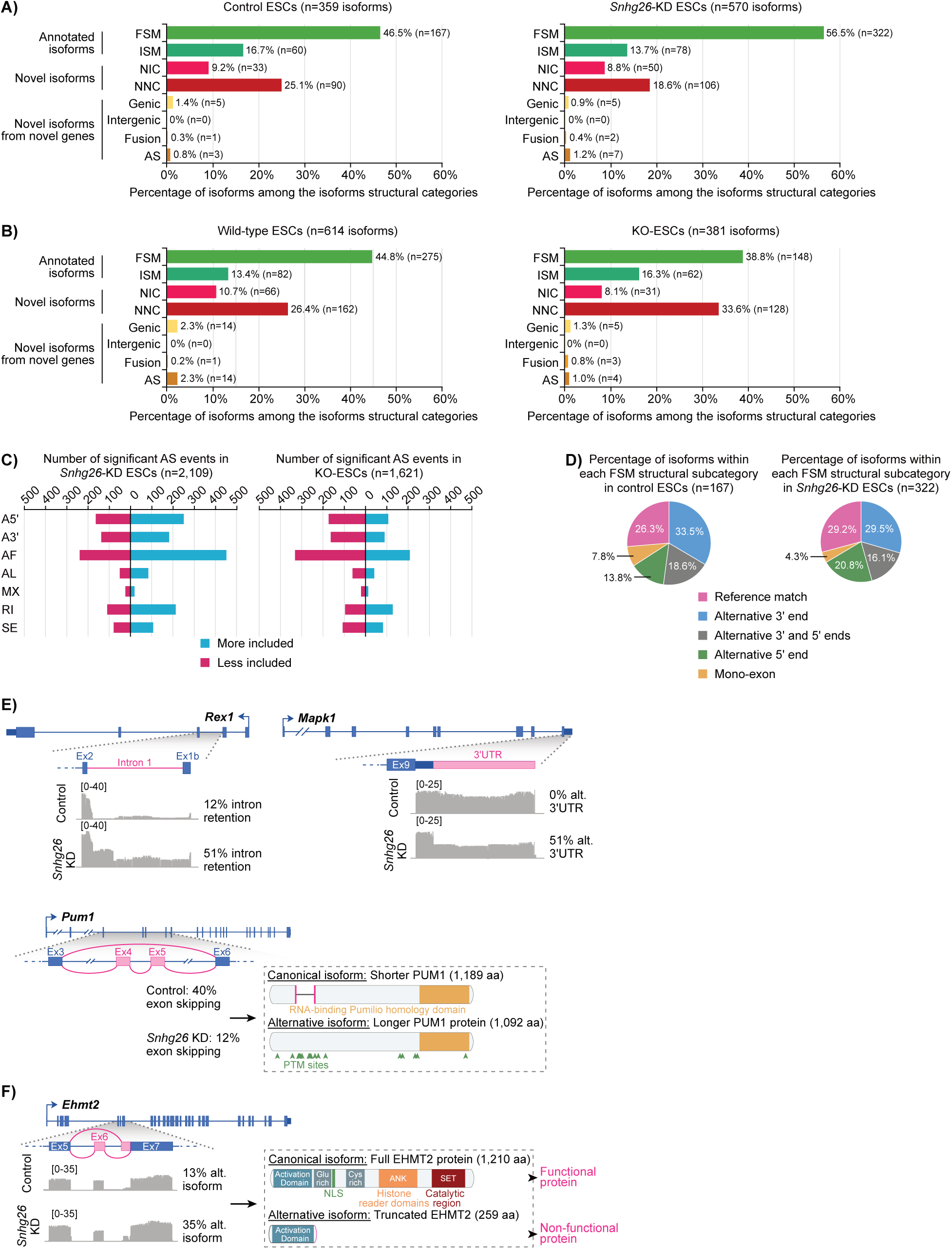
Loss of *Snhg26* impedes proper splicing of pluripotency and chromatin remodeling - associated transcripts. **A)** Bar graphs showing the proportion of alternatively spliced isoforms significantly enriched in control ESCs (left; n=4) and *Snhg26-*KD ESCs (right; n=2) within each isoform structural category. Significance based on a differential percentage spliced-in (ΔPSI) ≥ 0.1 (more included) or ΔPSI ≤ 0.1 (less included) and an adjusted p-value ≤ 0.05. FSM: Full-Splice Matches; ISM: Incomplete Splice Matches; NIC: Not-In-Catalogue; NNC: Novel-Not-in-Catalogue; AS: Antisense. **B)** Bar graphs showing the proportion of alternatively spliced isoforms significantly enriched in wild-type ESCs (left; n=2) and KO-ESCs (right; n=3) within each isoform structural category. Significance based on a ΔPSI ≥ 0.1 (more included) or ΔPSI ≤ 0.1 (less included) and an adjusted p-value ≤ 0.05. **C)** Bar graphs illustrating the number of significant alternative splicing events more (blue) or less (pink) included in *Snhg26-*KD ESCs (left) or KO-ESCs (right) compared to their control ESCs. Significance based on a ΔPSI ≥ 0.1 (more included) or ΔPSI ≤ 0.1 (less included) and an adjusted p-value ≤ 0.05. AS: Alternative Splicing; A5’: Alternative 5’ splice site; A3’: Alternative 3’ splice site; AF: Alternative First exon; AL: Alternative Last exon; MX: Mutually exclusive exons; RI: Retained Intron; SE: Skipped Exon. **D)** Pie charts representing the proportion of transcripts significantly enriched in control ESCs (left; n=4) and *Snhg26-*KD ESCs (right; n=2) within each FSM isoform structural subcategories. Significance based on a ΔPSI ≥ 0.1 (more included) or ΔPSI ≤ 0.1 (less included) and an adjusted p-value ≤ 0.05. **E)** Schematics highlighting the alternative splicing events (pink) within *Rex1* (left), *Mapk1* (right) and *Pum1* (bottom) gene loci following *Snhg26* KD for 5 days in ESCs, the associated lrRNAseq read coverage (grey tracks) and the resulting protein predictions. Ex: Exon; alt.: alternative; UTR: Untranslated Transcribed Region; aa: amino acid; PTM: Post-Translational Modification sites. Also see **Extended Data Fig. 3A**. **F)** Schematic highlighting the alternative splicing event (pink) within *Ehmt2* gene locus following *Snhg26* KD for 5 days in ESCs and the associated lrRNAseq read coverage (grey tracks). NLS: Nuclear Localization Sequence; ANK: Ankyrin domain; SET: Su(var)3-9, Enhancer-of-zeste and Trithorax domain. Also see **Extended Data Fig. 3B**.

To gain some insight into the functional consequences of such isoform diversity alteration, we searched for specific examples of alternative splicing. We found differential splicing, exon skipping, intron retention and alternative 3’UTR length in several of pluripotency- and chromatin remodeling-associated mRNAs. For instance, *Rex1*, a transcription factor that prevents differentiation and promotes proliferation of ESCs^41^ had significant intron retention (**Fig. 3E**). We found that after *Snhg26* KD, its first intron was prone to intron retention (12% in control cell lines vs 51% in *Snhg26-*KD ESCs), leading to an alternative isoform of *Rex1* that may be more prone to undergo nonsense-mediated decay^42^. Another alternatively spliced gene involved in maintenance of pluripotency was *Mapk1*. It is a serine/threonine kinase which acts as an essential component of the MAPK/ERK signaling pathway^43^. Following *Snhg26* KD in ESCs, 51% of *Mapk1* isoforms displayed a shorter 3’UTR (**Fig. 3E**). Other notable examples included *Stella*, which displayed half less proportion of isoforms with skipped exon 3 (20% in control cell lines vs 9% in *Snhg26-*KD ESCs) (**Extended Data Fig. 3A**). As *Stella* is important to maintain *naïve* pluripotency^44^, this alternative splicing event may explain the loss of *naïve* pluripotency observed when knocking-down *Snhg26*. Similarly to *Stella*, 5% of *n-Myc* isoforms, a gene involved in cell proliferation of pluripotent stem cells^45^, presented exon skipping in control ESCs. However, following *Snhg26* KD, we noted a complete disappearance of isoforms with skipped exon 2 (**Extended Data Fig. 3A**). A final example is *Pum1*, an RNA-binding protein involved in ESC maintenance and differentiation through degradation of specific mRNA targets^46^. In this case, *Snhg26* KD led to more inclusion of exons 4 and 5 (40% of exons skipping in control cell lines vs 12% exons skipping in *Snhg26-*KD ESCs) (**Fig. 3E**). Predictions of the protein domains informed that such AS event could lead to incorporation of phosphorylation sites involved in post-translational modifications. Phosphorylation of serine 714 is driven by growth factors and activates the binding of PUM1 to *Cdkn1b* mRNA, leading to degradation of the transcript and thus repression of *Cdkn1b* expression^47^. *Cdkn1b* was shown to control the G1 phase in ESCs, and its degradation promotes ESCs cycle progression^48^. Thus, *Snhg26* depletion in ESCs leads to alternative isoforms of genes involved in pluripotency and loss of *naïve* pluripotency.

Additionally, changes in isoform diversity did not only impact pluripotency-associated genes. Indeed, we also found alterations in the splicing of chromatin-remodeling genes, such as *Ehmt2*. The latter is a H3K9 methyltransferase involved in gene inhibition, particularly of transposable repeated sequences and of developmental genes, and is implicated in *de novo* methylation of pluripotency genes in pluripotent stem cells^49^. Following *Snhg26* KD, we found 2.5 more isoforms with skipping of exon 6 and exon 7 displaying a 5’ alternative splice site (13% in control cell lines vs 35% in *Snhg26-*KD ESCs) (**Fig. 3F**). This leads to an alternative isoform that loses both the ANK histone reader domain and the SET catalytic region of EHMT2, generating a truncated protein. Finally, *Brd8* is a bromodomain containing protein that recognizes acetylation marks on histones and anchors the acetyltransferase KAT5 to primed pluripotency-associated gene loci in order to maintain their expression^50^. In *Snhg26-*KD ESCs, *Brd8* exon 9 is skipped in 12% of isoforms, leading to an extended protein (**Extended Data Fig. 3B**).

Together, these results suggested that *Snhg26* lncRNA KD leads to an increase in novel isoforms, and that this alternative splicing landscape affects isoforms of pluripotency and chromatin remodeling-associated genes. Thus, *Snhg26* lncRNA may be involved in post-transcriptional regulation of these genes, which could explain the loss of the *naïve* pluripotent state of ESCs.

### 4. *Snhg26* interacts with transcripts of genes involved in embryonic pattern specification

To identify the mechanism of action of *Snhg26* lncRNA, we decided to identify its interactors in mouse ESCs. Because we previously showed that *Snhg26* lncRNA is mostly nuclear (Fort *et al.*, in preparation), and that its overexpression during reprogramming up-regulates chromatin-remodeling genes (**Extended Data Fig. 1D**), we first investigated whether *Snhg26* could interact with chromatin. To assess this, we first looked at published RADICLseq dataset^51^, which provides genome-wide RNA–chromatin interactions in crosslinked nuclei of ESCs. We found that *Snhg26* was not detected in proximity of chromatin genome-wide, except for its own locus (**Fig. 4A** and **Table S4**). To confirm this result, we performed chromatin isolation by RNA purification (ChIRP)^52^ in mouse ESCs using probes specific to *Snhg26.* We found no significant enrichment of *Snhg26* with chromatin using *Snhg26* probes or *LacZ* control probes, suggesting that *Snhg26* does not interact with DNA (**Fig. 4B** and **Table S4**). Finally, to exclude the possibility that the act of *Snhg26* transcription or that regulatory elements within its promoter could affect neighboring genes as it is sometimes the case for lncRNAs^53^, we analyzed gene expression 0.5 megabases up- and down-stream of *Snhg26* transcription start site following *Snhg26* KD. No significant change was observed in the 18 different genes that these regions encompass, suggesting that *Snhg26* does not act in *cis* on the genome to regulate gene expression (**Fig. 4C**).

**Figure 4:**
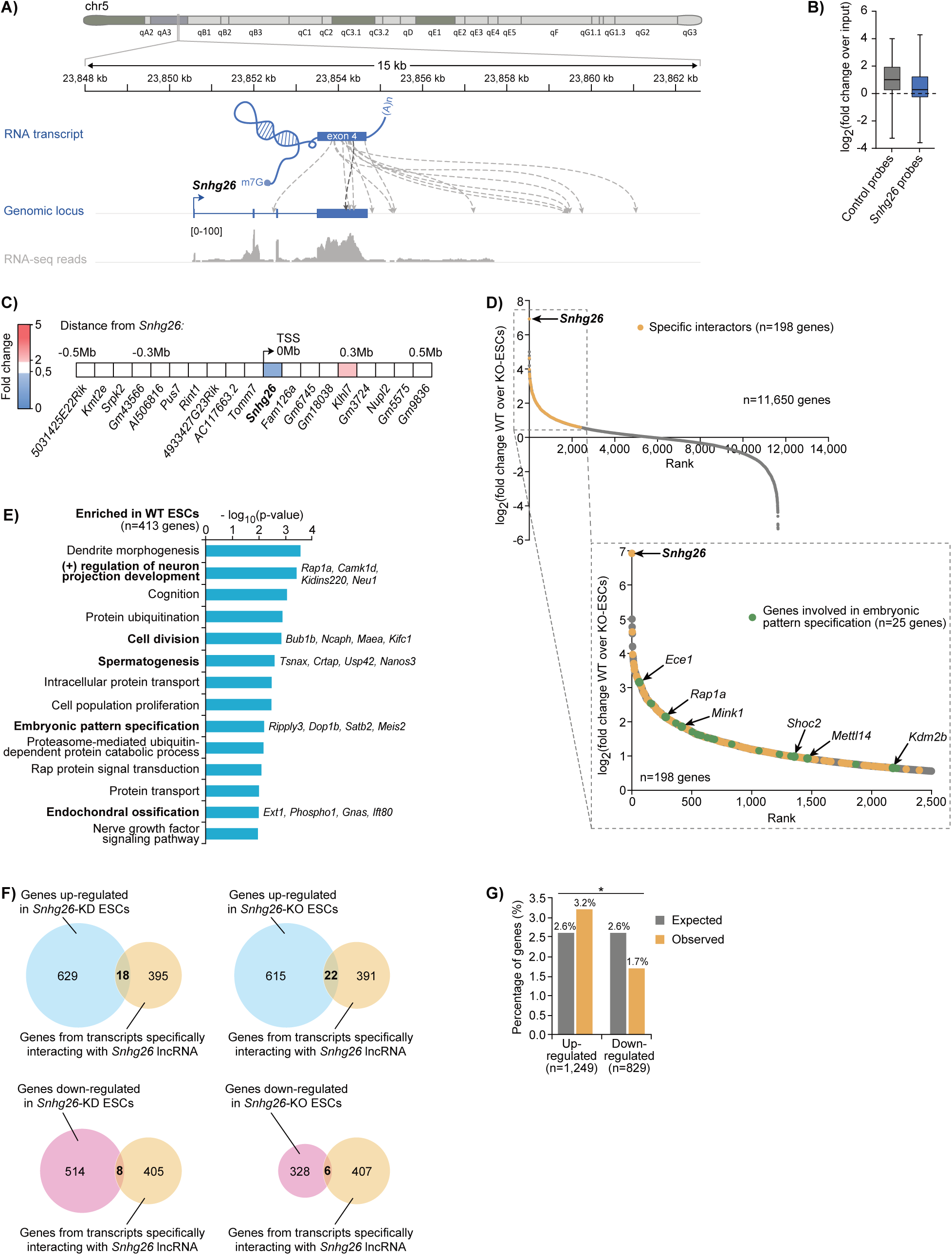
*Snhg26* interacts with transcripts of genes involved in embryonic pattern specification. **A)** Schematic representing all the *Snhg26* lncRNA interactions with chromatin detected in RADICLseq dataset^51^ from ESCs (upper track) as well as corresponding RNAseq read coverage (lower track). chr5: chromosome 5; kb: kilobases; m7G: 7-methylGuanylate cap; (A)n: poly(A) tail. **B)** Box plot representing the enrichment of *Snhg26* interactors detected by ChIRP-DNA over input using *Snhg26*-specific probes or control *LacZ* probes in ESCs. **C)** Schematic highlighting the expression fold change of genes located 0.5Mb up- and down-stream of *Snhg26* TSS in *Snhg26-*KD ESCs normalized to control ESCs. Mb: Megabases. **D)** Plot showing the ranking of genes of RNA transcripts (grey, n=11,650) detected by ChIRP-RNA in wild-type ESCs (n=5) compared to KO-ESCs (n=3) with their logged fold change. Transcripts significantly enriched (yellow, n=198) using *Snhg26*-specific even probes compared to control *GFP* probes are highlighted. Transcripts of genes involved in embryonic pattern specification (green, n=25) are also highlighted. Significance based on a fold change (wild-type ESCs over KO-ESCs) ≥ 1.5 and a p-value ≤ 0.05. Also see **Extended Data Fig. 3C**. **E)** Bar graph depicting the logged p-value of GO-terms of enriched biological processes for which genes are enriched in *Snhg26* ChIRP-RNA pulldowns with either *Snhg26-*specific even or odd probe sets (n=413). P-value < 0.025. **F)** Venn diagrams representing the overlap between the number of genes undergoing significant up-(blue) or down-(pink) regulation in *Snhg26-*KD ESCs (left) or KO-ESCs (right) and the number of genes which transcripts are significantly enriched in *Snhg26* pulldowns. **G)** Bar graph showing the expected (grey) and observed (yellow) percentages of genes that are common to *Snhg26* interactors and up-(n=1,249) or down-(n=829) regulated in *Snhg26-*KD ESCs or KO-ESCs. P-value was calculated by Chi-squared test. * < 0.05.

For the above reasons, we investigated possible *Snhg26* interactions with other RNA transcripts, as *Snhg26* may be involved in RNA post-transcriptional regulation. In order to do so, we modified the ChIRP protocol using a UV-aminomethyltrioxalen based crosslinking to specifically target RNA-RNA interactions^54,55^ (ChIRP-RNA) in mouse ESCs. As this type of experiment displays a lot of background noise signal coming from non-specific interactions, pulldown experiments were performed both in WT and KO-ESCs in order to identify more accurately interactions specific to *Snhg26* lncRNA. When comparing to non-specific probes targeting a *GFP* transcript (“control”), we were able to identify 413 genes specifically enriched in WT ESCs, with no significant enrichment in KO-ESCs, using two different sets of probes (**Fig. 4D**, **Extended Data Fig. 3C** and **Table S4**). GO-terms enrichment analysis for genes significantly pulled-down with either set of *Snhg26* probes included biological processes linked to embryonic pattern specification and tissue differentiation (e.g. “positive regulation of neuron projection development”, “spermatogenesis”, “endochondral ossification”) (**Fig. 4D-E**, **Extended Data Fig. 3C** and **Table S4**). When correlating the number of genes being up- and down-regulated in *Snhg26-*KD ESCs and KO-ESCs compared to control ESCs, within *Snhg26* interactors, we found only a few interactors are differentially expressed (**Fig. 4F**). However, interestingly, there is a significantly larger proportion of *Snhg26* interacting genes that are up-regulated than genes that are down-regulated when *Snhg26* is depleted or deleted (**Fig. 4G**). This suggests that *Snhg26* destabilizes some of its RNA interactors in ESCs.

### 5. *Snhg26* interacts with SINEB2 transposable sequences and affects their expression

In order to further characterize *Snhg26* mechanism of action, we screened its sequence in order to identify particular motifs or domains that could convey its function. Intriguingly, we found the presence of repetitive sequences derived from transposable elements. Indeed, studies examining functional motifs of lncRNAs have led to the RIDL hypothesis^15,16^, which postulates that lncRNA-embedded TEs could constitute functional units for these transcripts. On average, 69-83% of human lncRNAs span at least one TE, which is in stark contrast to protein-coding transcripts (6%)^16,18–21^. These estimations are lower in mice (33-68%), but they still demonstrate the high prevalence of TEs in lncRNAs^18,19,21^. Among human TE-containing lncRNAs, 19% of have at least 50% of their sequence derived from TEs, and those are often found within the last exon of the lncRNAs^19^. Strikingly, this was the case for the mature transcript of *Snhg26*, which presents four different TEs in its fourth exon: One SINEB1_Mus1, one SINEB2_Mm1a and two RLTR48_ERV1 sequences (670 nucleotides out of 1,385 total, i.e. 48.4%) (**Fig. 5A**). Because lncRNA-embedded TEs have been linked to several functions by allowing RNA-RNA duplex formation with other transcripts^17,22–25^, we decided to investigate whether *Snhg26* TEs could mediate any interaction with its target transcripts. First, we analyzed enrichment of the previously identified *Snhg26* RNA interactors for sequences derived from the different TE classes and orders. We found no enrichment for sequences derived from DNA transposons or LINE retrotransposons (**Fig. 5B**). However, SINE and to a lesser extent LTR sequences were enriched significantly compared to control probes (**Fig. 5B**). Because sequences derived from the SINEB1, SINEB2 and RLTR TEs were represented in *Snhg26* fourth exon, we further investigated if there was enrichment for these specific TE subfamilies (i.e. SINEB1_Mus1, SINEB2_Mm1a, RLTR48_ERV1) within *Snhg26* interactors. Interestingly, we observed a significant enrichment only for SINEB2_Mm1a TE-derived sequences, which corresponded to a 189 nucleotides-long sequence within the last 300 base pairs of *Snhg26* (**Fig. 5C**). This suggested that *Snhg26* interacts with hundreds of transcripts, with some possibly through complementary SINEB2 sequences, and that those interactions are lost upon *Snhg26* KO in ESCs. A surprising point was that most of these SINEB2_Mm1a were not embedded within transcripts but rather independent sequences, as suggested by the sharp edges of the SINE enrichment metaplots (**Fig. 5C**, bottom plots).

**Figure 5:**
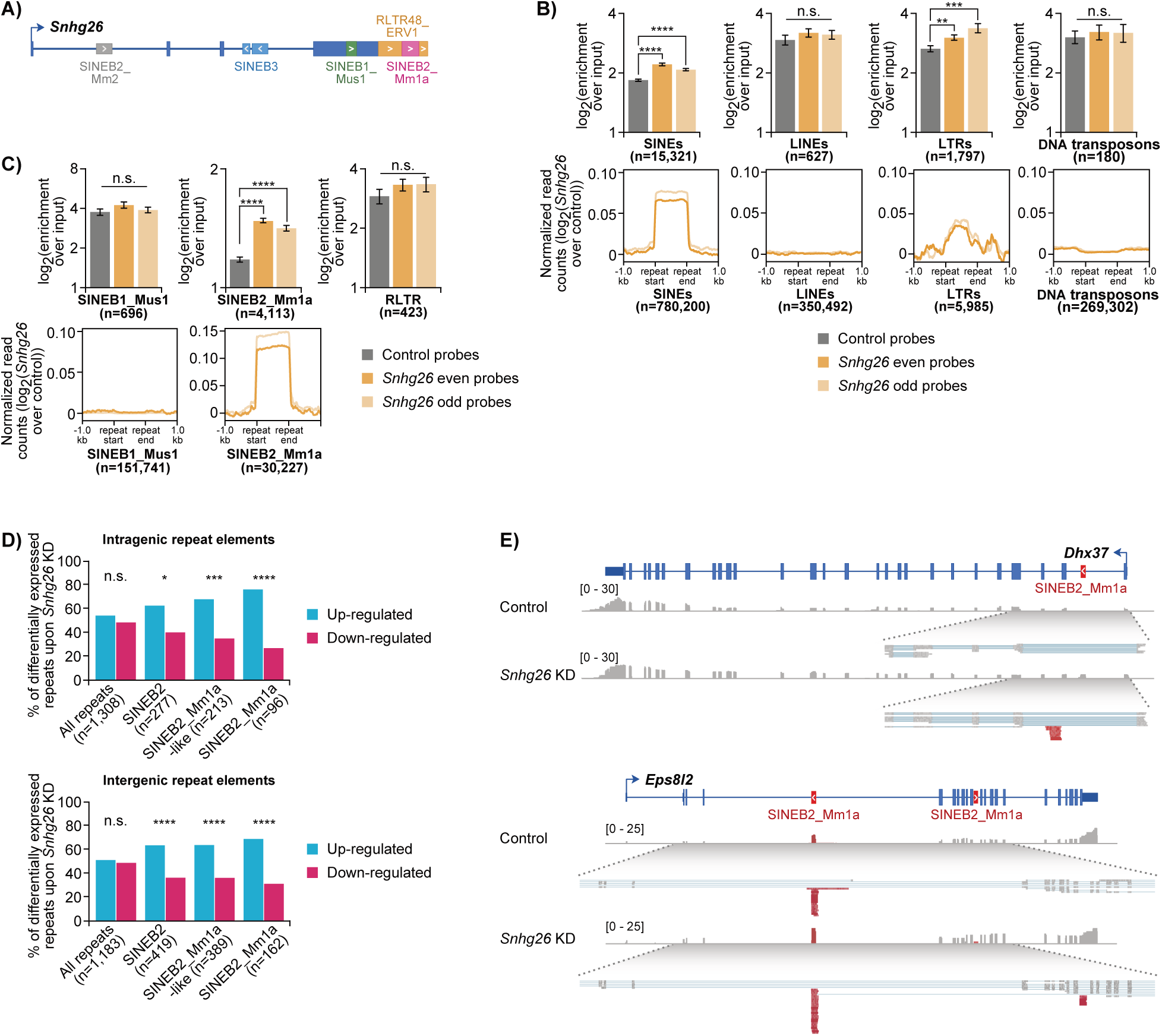
*Snhg26* interacts with SINEB2 transposable sequences and affects their expression. **A)** Schematic depicting *Snhg26* locus with its embedded transposable element-derived sequences. Arrows within colored boxes indicate presence of a transposable element and its orientation. **B)** Bar graphs and corresponding metaplots representing the logged average enrichment of the different transposable element super-families found within *Snhg26* interactors detected by ChIRP-RNA over input using *Snhg26*-specific probes or control *GFP* probes in ESCs. Error bars represent the SEM. P-values were calculated by two-tailed t-test (two samples with unequal variance). n.s. ≥ 0.05, ** < 0.01, *** < 0.001, **** < 0.0001. **C)** Bar graphs and corresponding metaplots representing the average enrichment of the different transposable element subfamilies found within *Snhg26* interactors detected by ChIRP-RNA over input using *Snhg26*-specific probes or control *GFP* probes in ESCs. Error bars represent the SEM. P-values were calculated by two-tailed t-test (two samples with unequal variance). n.s. ≥ 0.05, **** < 0.0001. **D)** Bar graphs showing the percentage of up-(blue) or down-(pink) regulated intragenic (top) and intergenic (bottom) transposable elements detected in ESCs after 5 days of *Snhg26* KD relative to control ESCs. P-values were calculated by Chi-squared test. n.s. ≥ 0.05, *** < 0.001, **** < 0.0001. **E)** Tracks showing the read coverage of *Dhx37* (top) and *Eps8l2* (bottom) gene loci containing SINEB2_Mma1 sequences (red) detected by lrRNAseq in control or *Snhg26-*KD ESCs.

To assess if this potential interaction could affect regulation of SINEB2 RNAs, we analyzed differential transposable elements expression in *Snhg26-*KD ESCs. Strikingly, we found that both intragenic and intergenic SINEB2 sequences were up-regulated in *Snhg26-*KD ESCs compared to WT ESCs (**Fig. 5D-E**). Moreover, this up-regulation was specific to SINEB2 TE family, and more particularly SINEB2_Mm1a repeats. This suggests that *Snhg26* regulates SINEB2_Mm1a repeat levels by direct interactions and this may lead to their degradation, as they are up-regulated when is *Snhg26* depleted.

### 6. The *Snhg26-*embedded SINEB2 sequence is essential for its function in pluripotency

We thus hypothesized that the SINEB2 element within *Snhg26* sequence may contribute to its lncRNA function in enhancing acquisition of the pluripotent state of iPSCs. To test this, we overexpressed a construct lacking only the SINEB2 element but leaving the rest of the sequence intact (*Snhg26*ΔSINEB2) along with OSKM transcription factors during reprogramming of NPCs into iPSCs. We measured the appearance of iPSCs by following the reactivation of a GFP reporter under the control of the *Oct4* gene using flow cytometry. Strikingly, replacing *Snhg26* by *Snhg26*ΔSINEB2 abrogated the effect of the lncRNA on reprogramming, as the percentage of GFP+ cells (i.e. iPSCs) at D16 went from 51.8% to 14.2%, which were almost similar levels as the control condition (28.3%) (**Fig. 6A**). To further validate this effect, we performed a limiting dilution assay to measure the frequency of iPSCs generated at D20 of OKS reprogramming with or without *Snhg26* SINEB2 element. We found again a lower percentage of iPSCs when *Snhg26*ΔSINEB2 was overexpressed (0.74%) than in the condition where the full-length *Snhg26* was expressed (1.47%), which was comparable to the control condition (0.55%) (**Fig. 6B**). Finally, to investigate the importance of *Snhg26* SINEB2 in the maintenance of ESCs, we generated a new KO-ESC line this time with a concomitant conditional expression of *Snhg26*ΔSINEB2. Interestingly, PCR amplification of *Snhg26* exon 4 followed by examination of migration on gel profiles revealed that overexpression of *Snhg26*ΔSINEB2 could not rescue establishment of homozygous KO clones (0% with control and *Snhg26*ΔSINEB2 transgenes), and only slightly increase the percentage of heterozygous clones (29.4% with control transgene vs 44.4% with *Snhg26*ΔSINEB2 transgene) (**Fig. 6C** and **Extended Data Fig. 3D**). Altogether, these results suggest that the SINEB2 element within *Snhg26* sequence is implicated in its function and in acquisition and maintenance of *naïve* pluripotency.

**Figure 6:**
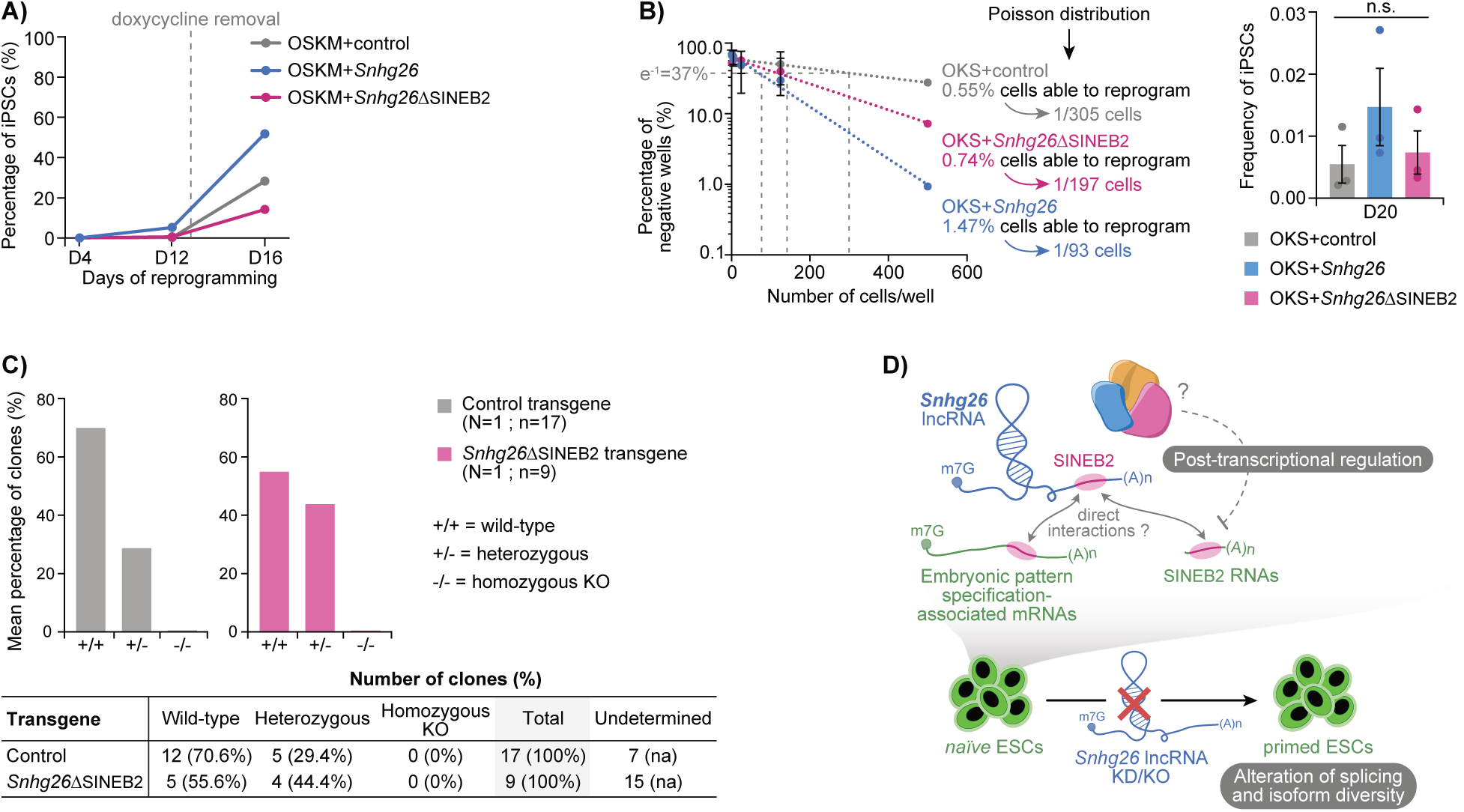
The *Snhg26*-embedded SINEB2 sequence is essential for its function in pluripotency. **A)** Graph representing the percentage of iPSCs in FBS-LIF + doxycycline medium, or in 2i-LIF medium after doxycycline removal at D13, monitored throughout reprogramming (n=1) using flow cytometry for OSKM+control (grey), OSKM+*Snhg26* (blue) and OSKM+ *Snhg26*ΔSINEB2 (pink) conditions. OSKM: *Oct4*, *Sox2*, *Klf4* and *c-Myc*. **B)** Graph (left) representing the mean percentage of GFP-negative wells (i.e. iPSCs-negative wells) for the different cell densities at D20 of reprogramming in 2i-LIF medium after doxycycline removal at D12 (n=3), allowing the use of Poisson distribution. Error bars represent the SEM. Bar graph (right) shows the corresponding average frequencies of iPSCs for OKS+control (grey), OKS+*Snhg26* (blue) and OKS+*Snhg26*ΔSINEB2 (pink) reprogramming conditions at D20 (n=3) of reprogramming. Error bars represent the SEM and each dot represents the replicates. P-values were calculated by one-tailed t-test with unequal variance. n.s. ≥ 0.05. **C)** Bar graphs (top) showing the mean percentage of ESC clones obtained with a wild-type (+/+), heterozygous (+/-) or homozygous KO (-/-) locus of *Snhg26* gene, for the conditions expressing a control (grey; n=17) or *Snhg26*ΔSINEB2 (pink; n=9) transgene. Table (bottom) indicates the raw number of clones (and the percentage they represent) obtained for N=1 experiment. Also see **Extended Data Fig. 3D**. **D)** Working model depicting how *Snhg26* lncRNA interactions with SINEB2 RNAs and genes involved in embryonic pattern specification through complementary SINEB2 derived sequences affects their post-transcriptional regulation in ESCs to maintain *naïve* pluripotency.

## Discussion

Across the literature, approximately fifty or so lncRNAs have been associated to mouse or human pluripotency^26,56–59^, but only a few are directly implicated in the regulatory network of the pluripotent cell states and embryonic development^60–65^. With our previous and current work, we add to this list by highlighting the role of *Snhg26* lncRNA in the maintenance of mouse ESCs and in enhancement of reprogramming.

We showed that *Snhg26* overexpression during induction of pluripotency activates earlier expression of genes involved in MET, signal transduction of pluripotency-related pathways and chromatin remodeling. As MET is promoted by BMP-SMAD signalling pathway, activation of SMAD and BMP signaling pathway-related genes as early as D2 of OKS+*Snhg26* reprogramming demonstrates an earlier commitment in MET^27,66^. Thus, overcoming of this reprogramming roadblock early shortens the initiation phase and explains the higher reprogramming yield when the lncRNA is expressed. Epigenetic modifications such as H3K9, H3K79, H3K36 and H3K27 methylation, as well as DNA methylation constitute another important barrier to reprogramming^67,68^. As chromatin remodeling genes were also enriched following *Snhg26* induction, it is possible that these epigenetic activators and repressors get regulated by *Snhg26* to favor the proper chromatin landscape modulation needed to achieve silencing of somatic genes and reactivation of pluripotency regulators that allows complete reprogramming to pluripotency. However, changes in the epigenetic marks of OKS+*Snhg26* reprogramming cells, as well as the specific mechanism of how *Snhg26* lncRNA could affect those changes, remain to be explored. Regarding *Snhg26* depletion and deletion, we showed that pluripotency- and chromatin remodeling-associated genes were affected as well. This reinforces the interest of investigating modifications in the chromatin landscape of ESCs when *Snhg26* expression is altered. Also, as GO-terms enrichment of DEGs in KO-ESCs revealed processes involved in embryonic pattern development, it would be of interest to further explore the incidence of *Snhg26* KD on mouse embryo development and the possibility of developmental defects. Finally, we noticed enrichment for genes involved in metabolism. Further investigations will be needed to determine if *Snhg26* has a role in metabolic regulation of ESCs and if this can affect loss of *naïve* pluripotency, as metabolic pathways have been shown to affect histone acetylation and gene expression involved in pluripotency maintenance^69–71^.

By taking advantage of lrRNAseq, we found that depletion and depletion of *Snhg26* lncRNA in ESCs alters isoform diversity, increasing the number of different transcripts in *Snhg26-*KD ESCs. However, an opposite effect was observed for KO-ESCs. This could be explained by the fact that despite *Snhg26* lncRNA expression levels being reduced by 80% after the KD (**Table S1**), the residual transcripts may exert a different function when lowered, especially if *Snhg26* function is dependent on the number of molecules in a cell. Despite KO-ESCs displaying less AS than control ESCs, intron retention was the only type of AS event enriched in both *Snhg26-*KD ESCs and KO-ESCs. Intron retention often results in mRNA decay and decreased protein translation, enabling rapid adaptation of gene expression without modulation of transcription^72^. Intron retention being involved in differentiation processes^73–75^, it would be of interest to characterize the genes undergoing intron retention and how it affects their expression. We also showed that following *Snhg26* KD, the higher number of isoforms was accompanied by more inclusion of AS events. Strikingly, several pluripotency and chromatin-remodeling genes underwent AS: A higher proportion of *Mapk1* transcripts displayed a shorter 3’UTR. As 3’UTRs are major regulators of mRNA stability and translation^76^, this alteration of splicing may have a profound impact on the *Mapk1* gene expression. According to protein predictions, alternative splicing of *Ehmt2* would lead to a non-functional protein. In the case of *Brd8,* it could affect the tertiary structure of BRD8 protein and lead to a more active isoform, increasing recruitment of KAT5 and further enhancing expression of primed pluripotency genes. Further investigation on how histone methylation and acetylation patterns are affected by depletion of *Snhg26* will thus be needed to measure further the impact of such observations. However, as genes involved in pluripotency and chromatin remodeling were enriched in the GO-terms of reprogramming, KD and KO of *Snhg26* lncRNA experiments, it would be interesting to evaluate the effect of those AS events on expression of their respective isoforms and if this impacts the global gene expression levels. Altogether, results from reprogramming and KD reinforced the idea that *Snhg26* lncRNA participates in the post-transcriptional regulation of the pluripotency regulatory network and the cooperation between chromatin remodelers to favor the pluripotent state. This was also supported by our finding that despite its nuclear localization, *Snhg26* does not bind to chromatin but to RNA transcripts of 413 different genes. Several of those are involved in regulation of embryonic pattern specification, such as *Klf7*, a transcription factor that binds *Nanog* regulatory regions^77^. Another target, *Satb2,* also associates to *Nanog* locus to facilitate its expression in ESCs^78^. Interestingly, *Snhg26* interacts as well with *Kdm2b* and *Hdac5* transcripts. The corresponding enzymes are implicated in regulation of the histone marks and epigenetic control of gene expression regulating the pluripotent state^79,80^. Thus, *Snhg26* may be involved in their post-transcriptional stabilization or alternative splicing.

In the pluripotency context or not, a major challenge in the field of lncRNAs is the systematic identification of their functional domains and interactome. While the development of techniques to grasp interactions between lncRNAs and chromatin ^81–83^, other RNA transcripts^84–86^ and proteins^87,88^ is continuously improving and allowing for deeper probing of lncRNA mechanisms of action, characterization of their functional domains remains challenging. The RIDL hypothesis has raised the concept of lncRNA-embedded TEs acting as functional domains^16,89^. So far, a few pioneering studies have shown examples of TEs acting as domains of lncRNAs to regulate their transcription^18,90,91^, post-transcriptional modifications^92^, localization^93^, stability^94^ and function^22,24,95,96^. A handful of studies have so far highlighted the role of SINEB2 TEs as functional domains. First, SINEB2s within certain lncRNAs have been shown to hybridize to complementary sequences on target mRNAs and to cause their degradation through Staufen1-mediated decay^24,25^. Second, several studies have shown that SINEB2 elements within antisense lncRNAs lead to activation of translation of the overlapping mRNA through a process called SINEUP^22,23^.

With our work, we identified within *Snhg26* lncRNA an embedded SINEB2 transposable element. Through enrichment analysis of TE-derived sequences present within *Snhg26* transcript interactors, we showed that this specific TE is present as well in several of *Snhg26* targets. We also found a strong up-regulation of intragenic and intergenic SINEB2 elements following depletion of *Snhg26* in ESCs, suggesting that the lncRNA regulates expression of SINEB2 TEs post-transcriptionally. This was striking as SINEB2 are repetitive elements that are present at 350,000 copies within the mouse genome (i.e. 2.39%)^97,98^, and their transcription is usually silenced by different epigenetic, piwiRNA, or transcription factor-mediated mechanisms in ESCs^98–101^. Indeed, in the pre-implantation blastocyst, DNA methylation levels silencing SINEs increase in the pluripotent epiblast cells and ESCs have global and retrotransposon DNA methylation levels similar to foetal somatic tissues^102–104^. This regulation is critical for maintaining genome integrity and preventing the potential harmful effects of transposon mobilization, which could lead to mutations or genomic instability^105–107^. We show that the SINEB2 within *Snhg26* sequence is necessary to its function in enhancing the efficiency of reprogramming towards pluripotent cells, and expression of *Snhg26* transcript lacking the SINEB2 sequence in *Snhg26*-KO cells could not lead to derivation of homozygous KO ESCs. This suggests that the SINEB2 element of *Snhg26* constitutes a functional domain mediating interactions with SINEB2 RNAs in ESCs in order to exert a post-transcriptional inhibiting effect on them.

Altogether, our work demonstrates the detailed consequences of *Snhg26* modulations on gene expression and isoform diversity during reprogramming to pluripotency and partial or total loss of *Snhg26* in ESCs. We bring to light its numerous interactions with multiple embryonic tissue specification - associated mRNAs, and also enrichment of interactors through a common embedded SINEB2-derived sequence. According to our working model, *Snhg26* and its SINEB2 functional domain are essential to maintain the pluripotent state of ESCs possibly through hybridization and destabilization of mRNAs involved in development and SINEB2 retrotransposable elements (**Fig. 6D**). Further work will be needed to establish whether this SINEB2 enables interactions that lead to post-transcriptional regulation of those targets. Thus, our study reiterates the importance of studying lncRNA functional domains and interactome to explore in detail its function and mechanism of action, and increases evidence for TE-derived functional domains carried through the RIDL hypothesis.

## Limitations of the study

The use of techniques relying on biotinylated antisense oligonucleotides to pull-down *Snhg26* lncRNA from ESC lysates may miss some of *Snhg26* interactors as it may not capture interactions that rely on intact subcellular architectures. Also, because of sequence constraints and repetitive elements, those *Snhg26*-specific probes could only be designed on a small portion of the lncRNA sequence (221 nucleotides, i.e. 16% of *Snhg26* lncRNA sequence). This may mask interactions mediated through this sequence. Moreover, direct interactions between *Snhg26* and its RNA targets have not been demonstrated to rely directly on the SINEB2 element. Finally, we cannot exclude that the other TEs or non-TE portions of *Snhg26* sequence also play a role in the biology of this lncRNA. However, importance of *Snhg26* SINEB2 sequence for its function is undeniable as its ablation abrogates improvement of reprogramming efficiency and derivation yield of *Snhg26* homozygous KO ESC clones.

## Authorship

S.H. conceived of and designed the study. V.F. and G.K. developed and performed the experiments and collected data. S.H., V.F. and G.K. designed and performed the computational and statistical analyses. V.F., G.K. and S.H. wrote, reviewed and edited the manuscript.

## Acknowledgments

We thank Valérie Watters for providing one of the EpiLC samples. This work was funded by the Canadian Institutes of Health Research (CIHR) to S.H (PJT-378019) and supported by the Fonds de recherche du Québec (FRQ) through the Research Centre Grant. V.F. is a recipient of a training award of the Fonds de recherche du Québec - Santé (FRQ-S). G.K. is a recipient of a training award of the Fonds de Recherche du Québec - Nature et Technologies (FRQ-NT). S.H. is a Junior 2 Research Scholar of the FRQS.

## Extended Data Figure legends

**Extended Data Figure 1:**
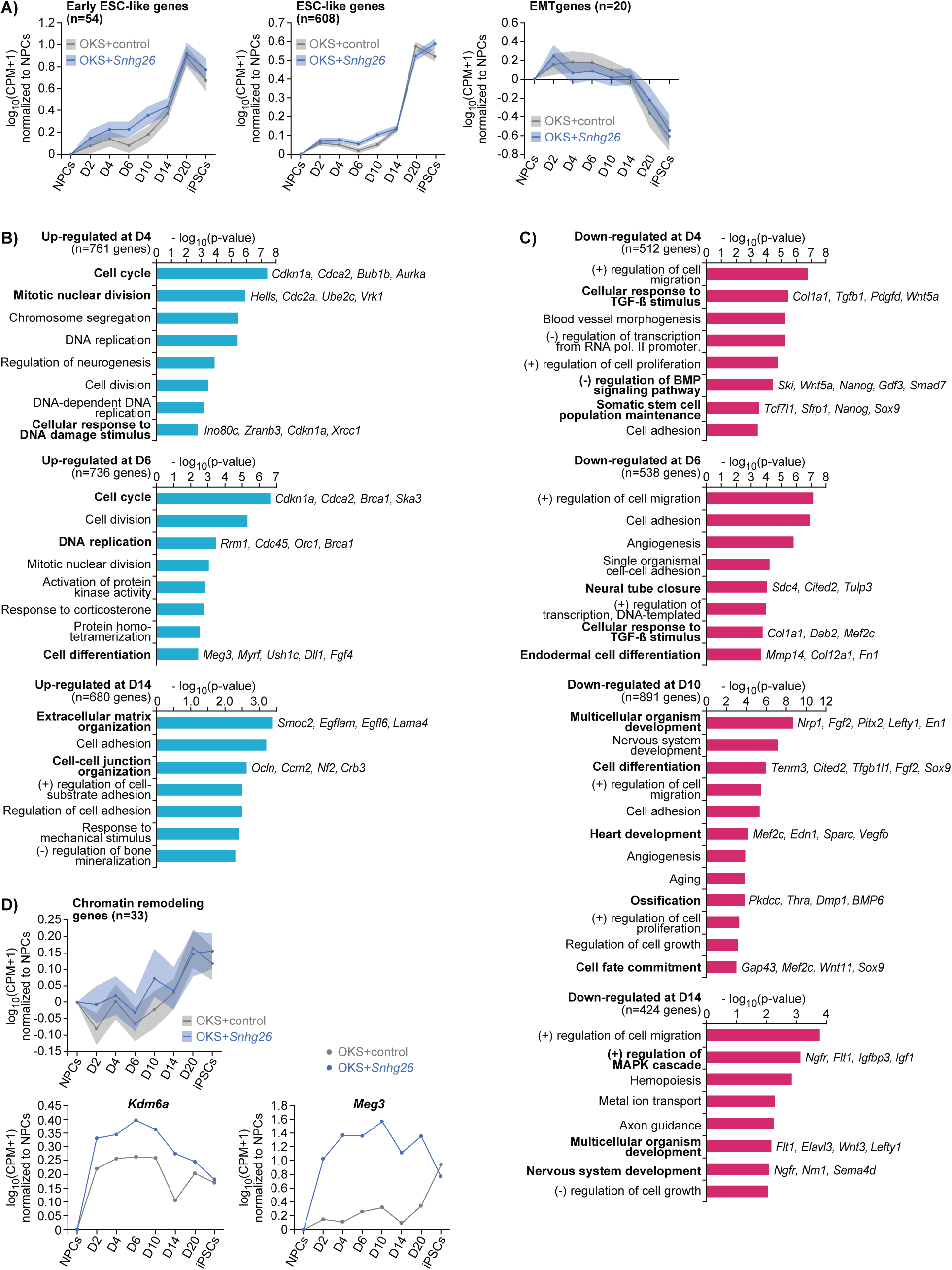
*Snhg26* lncRNA overexpression accelerates acquisition of pluripotency hallmarks early during reprogramming while loss of *Snhg26* promotes exit of the *naïve* pluripotency state, related to Figure 1. **A)** Plots depicting the logged fold change in expression in CPM of early ESC-like (n=54) and ESC-like (n=608) genes (gene lists from Hussein *et al.*, Nature 2014), or epithelial-to-mesenchymal transition (EMT) (n=20) genes in each timepoint of the OKS+control (grey) or OKS+*Snhg26* (blue) reprogramming, normalized to NPCs. Lines represent the average expression value at each timepoint and the shaded area represents the SEM. The data for OKS+control is the same as in Fort *et al.* (in preparation). **B)** Bar graphs depicting the logged p-value of GO-terms of enriched biological processes for which genes undergo up-regulation at D4 (n=761 genes), D6 (n=736 genes) and D14 (n=680 genes) of OKS+*Snhg26* reprogramming. P-value < 0.01. **C)** Bar graphs depicting the logged p-value of GO-terms of enriched biological processes for which genes undergo down-regulation at D4 (n=512 genes), D6 (n=538 genes), D10 (n= 891 genes) and D14 (n=424 genes) of OKS+*Snhg26* reprogramming. P-value < 0.01. **D)** Plot (top) depicting the logged fold change in expression in CPM of chromatin remodeling genes (n=33) in each timepoint of the OKS+control (grey) or OKS+*Snhg26* (blue) reprogramming, normalized to NPCs. Lines represent the average expression value at each timepoint and the shaded area represents the SEM. Graphs (bottom) showing the specific expression of *Kdm6a* and *Meg3* genes. The data for OKS+control is the same as in Fort *et al.* (in preparation).

**Extended Data Figure 2:**
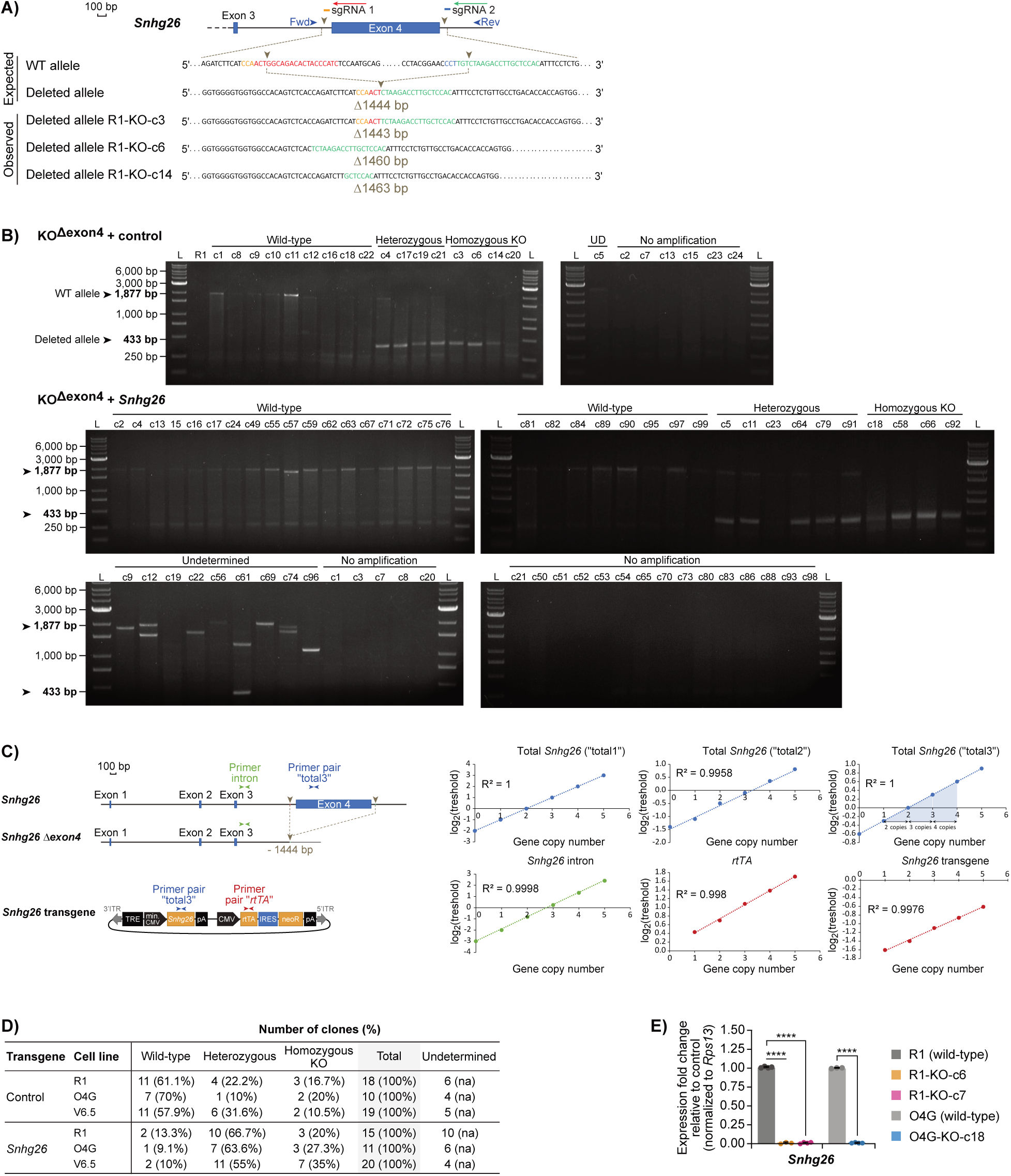
*Snhg26* lncRNA expression is essential for *naïve* pluripotency maintenance, related to Figure 2. **A)** Schematic of the *Snhg26* lncRNA locus and the result of exon 4 deletion following CRISPR-Cas9 technique on a wild-type allelic sequence. Blue arrows indicate the localization of the primer pair used for PCR screening. Orange and blue bars adjacent to sgRNAs represent the localization of their respective PAM motifs. Subjacent DNA sequences represent examples of exon 4 deletions in different R1-KO-ESC clones as obtained by Sanger sequencing of the PCR amplification products. Fwd: PCR screening Forward primer; Rev: PCR screening Reverse primer; WT: Wild-Type. **B)** Representative photographs of the migration profiles of the PCR amplification products obtained with the screening primers on R1 wild-type ESCs (n=1) or the different KO-ESC clones with control (n=24) or *Snhg26* (n=65) transgenes. DNA bands from the ladder corresponding to the associated length of the wild-type or deleted alleles are shown on the side. L: Ladder; UD: Undetermined. **C)** Graphs representing the linear regression curves allowing determination of the total vs transgenic number of copies of *Snhg26* allele in the KO-ESC clones by qPCR. “total1”, “total2” and “total3” correspond to three different primers. Same types of graphs are also showed for determining the number of copies of genomic intron 3 of *Snhg26* or the *rtTA* gene present on the vector carrying *Snhg26* transgene. R² is the coefficient of determination for each graph. **D)** Table showing the number of ESC clones (and the percentage they represent) obtained with a homozygous KO, heterozygous or wild-type locus of *Snhg26* gene, for the conditions expressing a control (n=47) or *Snhg26* (n=46) transgene for each cell line replicate (N=3 ESC lines). Some clones presented no amplification product by PCR and/or ambiguous copy number results by qPCR. These were not used in the analysis and designated as “undetermined”. na: non applicable. **E)** Bar graph representing the mean expression fold change of *Snhg26* in KO-ESC clones relative to their corresponding wild-type ESC lines (R1 and O4G), normalized to *Rps13* housekeeping gene. Error bars represent the STD. P-values were calculated by one-tailed t-test with unequal variance. **** < 0.0001.

**Extended Data Figure 3:**
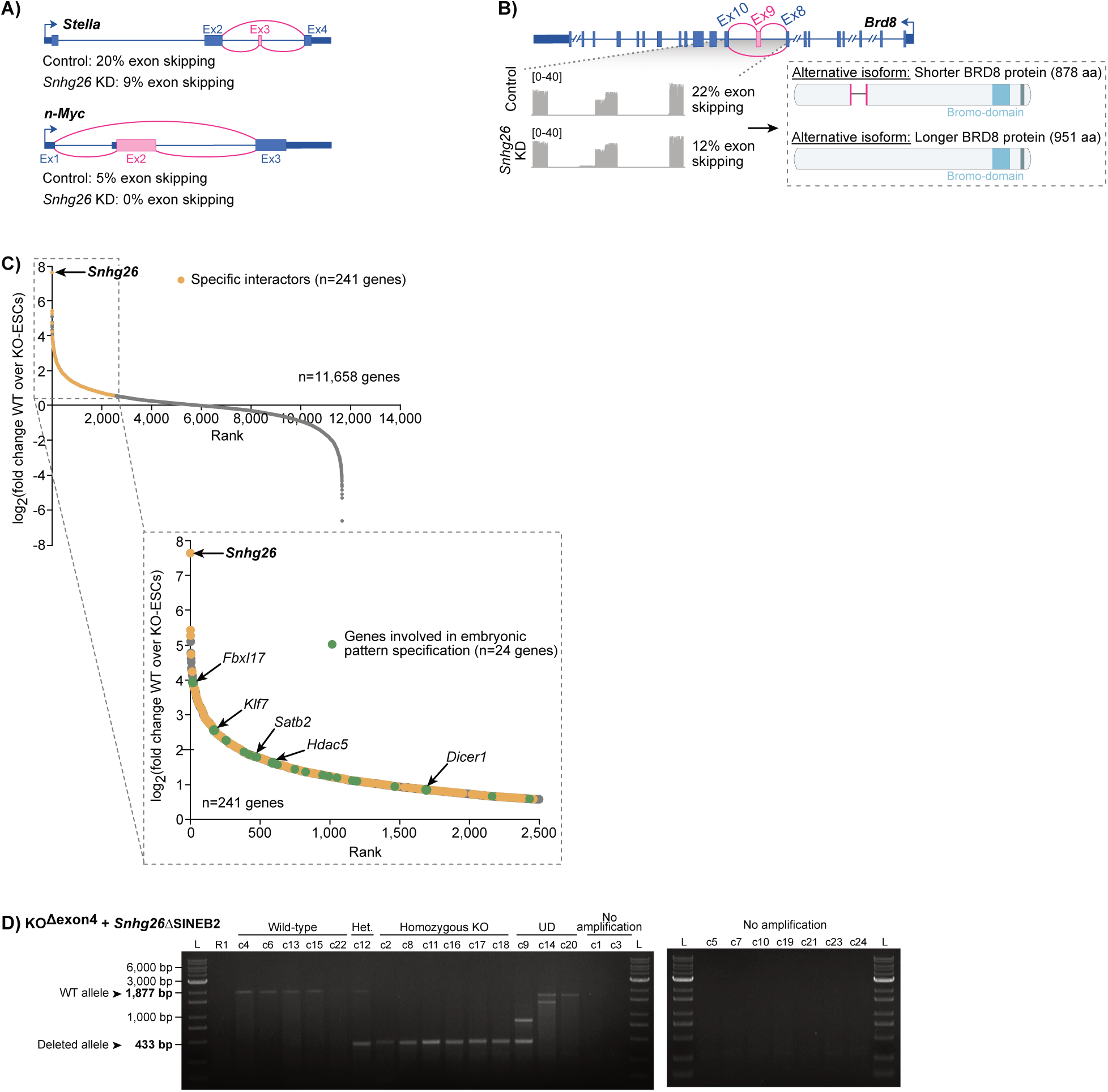
Loss of *Snhg26* impedes proper splicing of pluripotency and chromatin remodeling-associated transcripts, related to Figure 3, Figure 4 and Figure 6. **A)** Schematics highlighting the alternative splicing events (pink) within *Stella* and *n-Myc* gene loci following *Snhg26* KD for 5 days in ESCs. **B)** Schematic highlighting the alternative splicing event (pink) within *Brd8* gene locus following *Snhg26* KD for 5 days in ESCs, the associated lrRNAseq read coverage (grey tracks) and the resulting protein predictions. **C)** Plot showing the ranking of genes of RNA transcripts (grey, n=11,658) detected by ChIRP-RNA in wild-type ESCs (n=5) compared to KO-ESCs (n=3) with their logged fold change. Transcripts significantly enriched (yellow, n=241) using *Snhg26*-specific odd probes compared to control *GFP* probes are highlighted. Transcripts of genes involved in embryonic pattern specification (green, n=24) are also highlighted. Significance based on a fold change (wild-type ESCs over KO-ESCs) ≥ 1.5 and a p-value ≤ 0.05 **D)** Photographs of the migration profiles of the PCR amplification products obtained with the screening primers on R1 WT ESCs (n=1) or the different KO-ESC clones with *Snhg26*ΔSINEB2 transgene (n=24). DNA bands from the ladder corresponding to the associated length of the WT or deleted alleles are shown on the side. Het.: Heterozygous.

## Materials and methods

### Generation of the plasmid constructs

For cloning of the antisense of the *Luciferase* gene (control) and *Snhg26* transcripts, and for plasmid constructs used for reprogramming and knock-down experiments, please refer to our previous publication (Fort *et al.*, in preparation).

For the *Snhg26*ΔSINEB2 construct, specific primers were designed to PCR-amplify *Snhg26* without the SINEB2 element from the fourth exon (primer sequences are listed in **Table S2**). Proper sequence amplification was validated by Sanger sequencing and cloned into the PB-TetO-AIO destination vector using Gateway^TM^ cloning (ThermoFisher Scientific #11789020 & #11791020) strategy according to the manufacturer’s instructions. Inducible expression validated by RT-qPCR. Complete sequence of the resulting plasmid is available upon request.

### Reprogramming towards induced pluripotent stem cells

For generation and transfection of NPC lines for OKS and OSKM reprogramming, please refer to our previous publication (Fort *et al.*, in preparation). Stably transfected Oct4-GFP ESC-derived NPC cell lines were seeded at 3 x 10^3^ cells/cm^2^ on a 0.1% porcine gelatin (Sigma-Aldrich #G1890) coating in NPC medium supplemented with 2% FBS (ThermoFisher Scientific #12483020) to promote adhesion. The following day, medium was replaced with FBS-LIF supplemented with 1µg/mL doxycycline hyclate (Sigma-Aldrich #D9891) to induce overexpression of the reprogramming factors. Medium was refreshed every other day until D6. From D6 (unless mentioned otherwise), cells were maintained either in FBS-LIF + doxycycline or 2i-LIF medium and passaged every 2-4 days depending on colonies density. For passaging, cells were washed with DPBS (ThermoFisher Scientific #14190250) and treated with TrypLE^TM^ Express Enzyme (ThermoFisher Scientific #12604021) for 5 minutes. Then, TrypLE^TM^ Express Enzyme was neutralized using DMEM (high glucose, ThermoFisher Scientific #11965118) with 5% KnockOut^TM^ Serum Replacement (KSR, ThermoFisher Scientific #10828-028) and cells were passaged as single-cells (1/3-1/30 ratio depending on density). GFP+ cells were monitored using an Axio Vert.A1 (Zeiss) inverted fluorescence microscope with a 10X objective. Reprogramming cells were harvested using TrypLE^TM^ Express Enzyme for dissociation, washed with DPBS and pelleted for RNA extraction or resuspended in 500μL of DPBS for analysis using a BD Accuri^TM^ C6 Plus flow cytometer (BD Biosciences). iPSCs could readily be maintained in 2i-LIF medium after 20 days. All cell lines were negative for mycoplasma.

### Flow cytometry

Cells were harvested using TrypLE^TM^ Express Enzyme (ThermoFisher Scientific #12604021) for dissociation, washed with DPBS (ThermoFisher Scientific #14190250), resuspended in 500μL of DPBS and filtered through a 35μm cell strainer (Fisher Scientific #08-771-23). Samples were loaded into a BD Accuri^TM^ C6 Plus flow cytometer (BD Biosciences). Flow cytometry data were collected with the C6 Plus analysis software and analyzed with FlowJo^TM^ (v10.8.1). Cell debris were gated apart using an FSC-A x SSC-A plot. Then, singlets were gated using an FSC-A x FSC-H plot and the gating for fluorescent reporter-positive cells was determined using a reporter-negative control sample as well as separated samples each positive for the different reporter genes. The total number of live single cells, as well as the percentages of GFP+ cells and the volume used by the flow cytometer were collected directly from the .FCS files to calculate the density of GFP+ cells as follow: 𝑑𝑒𝑛𝑠𝑖𝑡𝑦 𝑜𝑓 𝐺𝐹𝑃 𝑐𝑒𝑙𝑙𝑠 = % 𝑜𝑓 𝑙𝑖𝑣𝑒 𝑠𝑖𝑛𝑔𝑙𝑒 𝐺𝐹𝑃 𝑐𝑒𝑙𝑙𝑠 × 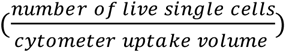. From this, the fraction of GFP+ cells was calculated as follow: % 𝐺𝐹𝑃 𝑐𝑒𝑙𝑙𝑠 = 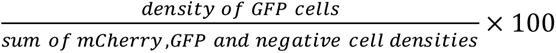. The values of 1 experiment were represented using

### Limiting dilution assays

Oct4-GFP ESC-derived NPC cell lines stably transfected with OKS reprogramming factors and control, *Snhg26* or *Snhg26*ΔSINEB2 transgenes were seeded at 1, 5, 25, 125, or 500 cells per well of a Falcon ^TM^ 96-well Flat-Bottom microplate (Fisher Scientific #08-772-2C) on a 0.1% porcine gelatin (Sigma-Aldrich #G1890) coating in NPC medium. The following day, medium was replaced with FBS-LIF medium supplemented with 1µg/mL doxycycline hyclate (Sigma-Aldrich #D9891) to induce the reprogramming into iPSCs. Medium was refreshed every other day for two weeks and switched at D12 to 2i-LIF medium until reprogramming completion. The number of wells negative for GFP+ cells were quantified manually using an Axio Vert.A1 (Zeiss) inverted fluorescence microscope with a 10X objective after 20 days. In Microsoft Excel, the average percentage of negative wells was plotted on a semi-logarithmic graph in function of the number of cells seeded as an exponential line, with error bars representing the SEM. The Poisson distribution allowed to determine the number of cells (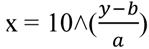, where y = 37, a = slope and b = y intercept) that must be seeded to have 37% of negative wells. The frequency (1/x) and percentage (1/x *100) of cells able to reach the iPSC stage were calculated based on three independent experiments (n=3). Frequencies of reprogramming cells and iPSCs were represented with SEM using GraphPad Prism (v7.04). P-values between the frequencies were calculated by one-tailed t-test (two samples with unequal variance). n.s. ≥ 0.05.

### Mouse ESC culture

ESC lines, including R1 (RRID: CVCL_2167), Oct4-GFP (R1 ESCs harboring a GFP reporter gene under the control of the *Oct4* gene promoter^108^) (both gifted by Dr Andras Nagy, University of Toronto), and V6.5 (gifted by Dr Steve Bilodeau, Laval University) were routinely maintained in a humidified incubator at 37°C under 5% CO2. Cells were propagated in 2i-LIF medium on 0.1% porcine gelatin (Sigma-Aldrich #G1890). 2i-LIF medium consists of Leukemia Inhibitory Factor (LIF, home-made from pGEX-2T-MLIF^109^) at 20ng/mL, 3μM CHIR-99021 (Selleck Chemicals #S2924) and 1μM PD0325901 (Selleck Chemicals #S1036) in N2B27 media. N2B27 media consists of a 1:1 mixture of DMEM/F-12 media mix (ThermoFisher Scientific #11330057) supplemented with 1X N-2 supplement (ThermoFisher Scientific #17502048) and 20μg/mL human recombinant Insulin (ThermoFisher Scientific #12585014), and Neurobasal^TM^ media (ThermoFisher Scientific #21103049) supplemented with 1X B-27^TM^ supplement (ThermoFisher Scientific #17504044), 1mM sodium pyruvate (ThermoFisher Scientific #11360070), 2mM L-glutamine (ThermoFisher Scientific #35050061), 1X MEM Non-Essential Amino Acid solution (ThermoFisher Scientific #11140050) and 10^-4^M 2-Mercaptoethanol (ThermoFisher Scientific #21985023). Medium was refreshed every other day and ESCs were passaged every 2-3 days. For passaging, cells were washed with DPBS (ThermoFisher Scientific #14190250) and treated with TrypLE^TM^ Express Enzyme (ThermoFisher Scientific #12604021) for 5 minutes. Then, TrypLE^TM^ Express Enzyme was neutralized using DMEM (high glucose, ThermoFisher Scientific #11965118) with 5% KSR (ThermoFisher Scientific #10828-028) and cells were passaged as single-cells (usually 1/10 ratio). All cell lines were negative for mycoplasma.

### *Snhg26* KO in mouse ESCs

KO-ESC lines (including R1, Oct4-GFP and V6.5) were established as described in Ran *et al*., Nature Protocols 2013 using two sgRNAs to remove the full fourth exon of *Snhg26* in ESC lines expressing either the antisense of the *Luciferase* gene (control), full-length *Snhg26* or *Snhg26ΔSINEB2* transcripts. Wild-type ESCs with no transgene were also used as controls. Single guide RNAs (sgRNAs) were designed using the CRISPR 10K Tracks of the UCSC genome browser^110^ and ordered from IDT^TM^. 0.350ng of annealed sgRNA pairs were cloned in 50ng of the pCCC backbone^111^ (gifted by Dr Francis Lynn, University of British Columbia) by BbsI HF® digestion (NEB #R3539) followed by ligation using a T4 DNA ligase (ThermoFisher Scientific #EL0016) according to the manufacturer’s instructions (sgRNA sequences and primers to clone transgenes are listed in **Table S2**). The sequences of all entry clone inserts were verified by Sanger sequencing and inducible expression validated by RT-qPCR. Complete sequences of all plasmids are available upon request.

For targeting with the CRISPR-Cas9 system, ESC lines cultured in a humidified incubator at 37°C under 5% CO2 were seeded at a density of 13 x 10^3^ cells/cm^2^ on plates coated with 0.1% porcine gelatin (Sigma-Aldrich #G1890) in 2i-LIF medium. The following day, cells were transfected with 50ng of hyPBase transposase (gifted by Dr Andras Nagy, University of Toronto), 800ng of each sgRNA constructs and 200ng of PB-TetO-AIO constructs carrying the control, *Snhg26* or *Snhg26*ΔSINEB2 constructs using Lipofectamine^TM^ Stem Transfection Reagent (ThermoFisher Scientific #STEM00015) according to the manufacturer’s instructions. Cells were then selected with 1μg/mL puromycin (Bio Basic #PJ593) for 4 days or 400μg/mL G-418 sulfate (Wisent #400-130-IG) for 6 days. Medium was supplemented with 100ng/mL doxycycline hyclate (Sigma-Aldrich #D9891) to induce expression of the transgenes. Five days post-selection, cells were dissociated as single-cells, counted and seeded at one cell per well of a Falcon ^TM^ 96-well Flat-Bottom Microplate (Fisher Scientific #08-772-2C) with a multichannel pipettor for colony isolation. Medium was replaced every other day and wells containing single colonies were selected by eye using an Axio Vert.A1 (Zeiss) inverted microscope with a 10X objective. One week post-seeding, clones were split, expanded and frozen prior to screening and further characterization. All cell lines were maintained in 2i-LIF medium supplemented with 100ng/mL doxycycline hyclate on 0.1% porcine gelatin, in a humidified incubator at 37°C under 5% CO2 and were negative for mycoplasma.

### KO-ESC clones screening and validation

During clone amplification, genomic DNA (gDNA) was extracted using gSYNC^TM^ DNA Extraction Kit (Geneaid #GS100) following the manufacturer’s instructions, with 100μg/mL RNaseA treatment (ThermoFisher Scientific #EN0531) 20 minutes at room temperature. DNA was quantified using a NanoDrop^TM^ 2000 Spectrophotometer (ThermoFisher #ND-2000C) and 300ng of DNA were loaded onto a 1% Agarose A (Bio Basic #D0012) gel to verify the DNA integrity and purity.

Assessment of ESC clones genotype regarding *Snhg26* locus was performed by PCR on 2ng of gDNA using the Deoxynucleotide (dNTP) Solution Set (NEB #N0446S) and the Taq DNA polymerase (NEB #M0267) according to the manufacturer’s instructions. 15-25μL of PCR-amplified product was loaded on a 1% agarose gel with 5μL of the GD 1kb plus DNA Ladder RTU (FroggaBio #DM015-R500F) and revealed using an Azure 200 gel image (Azure Biosystems). PCR-amplification of *Snhg26* fourth exon screening followed by Sanger sequencing validation was used to assess the copy number of *Snhg26* allele in each clone (genotyping primer sequences are described in **Table S2**). For validation, qPCRs were performed using 1ng of gDNA per clone (with a minimum of six technical replicates) and SYBR^TM^ Select Master Mix (ThermoFisher Scientific #4472908) on a CFX384 Touch real-time PCR detection system (Bio-Rad). Raw data was retrieved using the Bio-Rad CFX Maestro (v5.3.022.1030) software.

Allele copy number was analyzed on Microsoft Excel using the following method: *Sox1* locus (2 copies in any edited or wild-type clone) was used as control for normalizing the gDNA quantity (ng) for each clone. From this, a linear regression curve was generated to extrapolate the Cq values corresponding to the total copy number of total *Snhg26* and control/*Snhg26-*transgene copy number. From this, the number of endogenous copies of *Snhg26* was deducted by subtraction (total - transgene copy number). Wild-type ESCs with no transgenes as well as primers detecting the intron 3 of *Snhg26* were used as control for *Snhg26* locus (2 copies) (qPCR primer sequences are listed in **Table S2**). The percentage of wild-type, heterozygous or homozygous KO clones were calculated over the total number of validated clones (i.e. belonging to one of the aforementioned categories). The average percentage of independent experiments on three different cell lines (R1, Oct4-GFP and V6.5 cells) were used and represented with SEM using GraphPad Prism (v10.0). P-values were calculated by one-tailed t-test (two samples with equal variance). In total, 47 (control transgene), 46 (*Snhg26* transgene) ESC clones across all cell lines, and 9 (*Snhg26*ΔSINEB2 transgene) ESC clones for R1 cell line, were used for the analysis. Some clones presented no amplification product by PCR and/or ambiguous copy number results by qPCR. These were not used in the analysis and designated as “undetermined”.

### EpiLC differentiation

ESCs were seeded at 3 x 10^3^ cells/cm^2^ on Geltrex^TM^-coated plates (hESC-qualified, growth factor-reduced, ThermoFisher Scientific, #A1413302) in EpiLC medium. EpiLC medium consists of 20ng/mL recombinant Activin A (Peprotech #120-14E), 12ng/mL human recombinant FGF-2 (Peprotech #100-18B) and 1% KSR (ThermoFisher Scientific #10828-028) in N2B27 media. Half-medium change was performed daily. Differentiated cells were harvested D1, D2 and D3 using TrypLE^TM^ Express Enzyme for dissociation, washed with DPBS and pelleted for RNA extraction. All cell lines were negative for mycoplasma.

### RNA extraction and reverse transcription-qPCR

Cell lysates were homogenized with QIAshredder columns (Qiagen #79656) and total RNA was extracted using RNeasy Kit (Qiagen #74106) according to the manufacturer’s instructions. RNA was quantified using a NanoDrop^TM^ 2000 Spectrophotometer (ThermoFisher #ND-2000C) and then DNase-treated using DNaseI (ThermoFisher Scientific #EN0521) according to the manufacturer’s instructions. 300ng of DNase-treated and non-treated RNA were loaded onto a 1% Agarose A (Bio Basic #D0012) gel to verify the RNA integrity and purity. cDNA was synthesized from 500ng of RNA by oligo-dT and random priming with iScript^TM^ gDNA Clear cDNA Synthesis Kit (Bio-Rad #1725035), according to the manufacturer’s instructions. cDNA was diluted ten times and qPCRs were performed using SYBR^TM^ Select Master Mix (ThermoFisher Scientific #4472908) on a CFX384 Touch real-time PCR detection system (Bio-Rad). Raw data was retrieved using the Bio-Rad CFX Maestro (v5.3.022.1030) software. Relative gene quantification was analyzed based on the 2ΔΔCt method by applying the comparative cycle threshold (Ct) method on Microsoft Excel. Relative expression levels were normalized to *Rps13* housekeeping gene (qPCR primer sequences are listed in **Table S2**).

### Library preparation and long-read RNA-sequencing

Cell lysates were homogenized with QIAshredder columns (Qiagen #79656) and total RNA was extracted using RNeasy Kit (Qiagen #74106) according to the manufacturer’s instructions (RNAseq samples are listed in **Table S1**). RNA was quantified using a NanoDrop^TM^ 2000 Spectrophotometer (ThermoFisher #ND-2000C) and then DNase-treated using TURBO^TM^ DNase (ThermoFisher Scientific #AM2238) according to the manufacturer’s instructions. 300ng of DNase-treated and non-treated RNA were loaded onto a 1% Agarose A (Bio Basic #D0012) gel to verify the RNA integrity and purity. RNA was also quantified using the Qubit™ RNA Broad Range Kit (ThermoFisher Scientific #Q10210) on Qubit ™ 4 Fluorometer (ThermoFisher Scientific #Q33238) according to the manufacturer’s instructions. RNA quality was further verified using an RNA 6000 Nano Kit (Agilent Technologies #5067-1511) on a 2100 Bioanalyzer instrument (Agilent Technologies #G2939B) according to the manufacturer’s instructions. Samples all had RNA integrity numbers (RINs) of 7.8 or higher, except for mESCs_R1 (6.9).

For samples sequenced on R9 Flow Cells, polyadenylated RNA was enriched from 5-20μg of total RNA using NEBNext® Poly(A) mRNA Magnetic Isolation Module (NEB #E7490) according to the manufacturer’s instructions. Poly(A)+-enriched RNA was quantified using the Qubit ™ RNA High Sensitivity Kit (ThermoFisher Scientific #Q32852) on Qubit ™ 4 Fluorometer according to the manufacturer’s instructions and poly(A)+-enriched RNA smear quality was verified using an RNA 6000 Pico Kit (Agilent Technologies #5067-1513) on a 2100 Bioanalyzer instrument according to the manufacturer’s instructions. Libraries were generated using 55-100ng of poly(A)+ RNA and a Direct cDNA Sequencing Kit (Oxford Nanopore Technologies (ONT) #SQK-DCS109), with each sample barcoded with the Native Barcoding Expansion (ONT #EXP-NBD104), according to the manufacturer’s protocol: “Direct cDNA Native Barcoding (SQK-DCS109 with EXP-NBD104 and EXP-NBD114)” version DCB_9091_v109_revF_04Feb2019. Barcoded samples were pooled, loaded on a MinION sequencer (ONT) using eight R9.4.1 Flow Cells (ONT #FLO-MIN106D), and sequenced using the MinKNOW^TM^ (v19) software. Purification of DNA between each step was performed with KAPA HyperPure Beads (Roche #08963835001). R9.4.1 Flow cells were washed and reloaded with the Flow Cell Wash Kit (ONT #EXP-WSH003) to increase sequencing depth obtained per Flow cell.

For samples sequenced on R10 Flow Cells, Ligation sequencing amplicons - Native Barcoding Kit 24 V14 (ONT #SQK-NBD114.24) was used with 50ng of total RNA. Each sample was assigned a specific barcode. Briefly, an oligo-dT VN Primer (VNP, sequence: 5’-5phos/ACTTGCCTGTCGCTCTATCTTCTTTTTTTTTTTTTTTTTTTTVN-3’, where V=A, C, or G, and N=A, C, G, or T) was used to reverse-transcribe poly(A) RNA into cDNA with the Maxima H Minus Reverse Transcriptase (RT) (ThermoFisher Scientific #EP0751). A Strand Switching Primer (SSP, sequence: 5’-TTTCTGTTGGTGCTGATATTGCTmGmGmG-3’, where mG = 2’ O-Methyl RNA bases) was used for strand switching during the RT reaction. Libraries were PCR-amplified for 14 PCR cycles with ONT’s cDNA Primers (cPRM) (Forward primer sequence: 5’-ATCGCCTACCGTGACAAGAAAGTTGTCGGTGTCTTTGTGACTTGCCTGTCGCTCTATCTTC-3’, Reverse primer sequence: 5’-ATCGCCTACCGTGACAAGAAAGTTGTCGGTGTCTTTGTGTTTCTGTTGGTGCTGATATTGC-3’) and the LongAmp® Taq 2X Master Mix (NEB #M0287), following ONT’s SQK-PCS109 kit recommendations for PCR incubation times and temperatures. cDNA was quantified using the Qubit ™ 1X dsDNA High Sensitivity Kit (ThermoFisher Scientific #Q33230) on Qubit ™ 4 Fluorometer according to the manufacturer’s instructions and cDNA smear quality was verified using a DNA 12000 Kit (Agilent Technologies #5067-1508) on a 2100 Bioanalyzer instrument (Agilent Technologies #G2939B) according to the manufacturer’s instructions. The blunt DNA ends were prepared using the NEBNext® Ultra^TM^ II End Repair/dA-Tailing module (NEB #E7546), adding a 3’ dA tail and phosphorylating the 5’ end. Barcodes were ligated to the DNA fragments with a Blunt/TA ligase (NEB #M0367), barcoded samples were pooled and motor protein adapters were finally ligated with the NEBNext® Quick Ligation Module (NEB #E6056). Purification of DNA between each step was performed with KAPA HyperPure Beads (Roche #08963835001). Library was loaded on a PromethION 2 Solo (ONT) sequencer using two R10.4.1 Flow Cells (ONT #FLO-PRO114M) and sequenced using the MinKNOW^TM^ (v23.07.15) software. LrRNAseq datasets of reprogramming samples for OKS+control were previously analyzed in Fort *et al.* (in preparation). Raw fast5 files from sequencing with R9.4.1 Flow cells were converted to the more recent pod5 format using the pod5-file-format software from ONT’s “nanoporetech” GitHub page (https://github.com/nanoporetech/pod5-file-format). pod5 files were then basecalled using Guppy (v6.5.7) using super accurate settings with the guppy_basecaller command and the dna_r9.4.1_450bps_sup.cfg configuration file. Barcodes were detected and reads were demultiplexed with the guppy_barcoder command using the –barcode_kits “EXP-NBD104 EXP-NBD114” argument. Raw pod5 files from sequencing with R10.4.1 Flow cells were basecalled using Guppy (v6.5.7) using super accurate settings with the guppy_basecaller command and the dna_r10.4.1_e8.2_400bps_sup.cfg configuration file. Barcodes were detected and reads were demultiplexed with the guppy_barcoder command using the – barcode_kits “SQK-NBD114-24” argument. The resulting FASTQ reads were aligned to the GRCm38/mm10 mouse reference genome with Minimap2^112^ (v2.24-r1122) using the following parameters: -aLx splice –cs = long. Aligned reads were quality-controlled with the NanoPlot software^113^ (v1.43.2).

### LrRNAseq data analysis of gene expression

Raw read counts were obtained with the featureCounts tool^114^ from the Subread package (v2.0.6), using the exon counting mode. EdgeR R- package^115^ (v3.12.1) was then used to normalize the data, calculate RNA abundance at the gene level (as counts per million reads (CPM)), and perform statistical analysis. Briefly, a common biological coefficient of variation (BCV) and dispersion (variance) were estimated based on a negative binomial distribution model. This estimated dispersion value was incorporated into the final EdgeR analysis for differential gene expression, and the generalized linear model (GLM) likelihood ratio test was used for statistics, as described in EdgeR user guide. Differential gene expression was established as genes with a fold change ≥ 1.25 (up-regulation) or ≤ 0.8 (down-regulation) and and p-value ≤ 0.05 (calculated by 1-tailed t-test with unequal variance). Multidimensional scaling analysis was generated using “plotMDS” function within the EdgeR R-package.

Gene expression plots were generated using Microsoft Excel by averaging the log10(CPM+1) – log10(CPM_NPCs_+1) of different gene lists and representing the SEM as a shaded contour (gene lists are indexed in **Table S1**). GO-terms enrichment and KEGG pathway were assessed using the Database for Annotation, Visualization and Integrated Discovery (DAVID) (v2022q3)^116^. Biological Processes (BP) and KEGG pathway annotations were analyzed with a cut-off of p-value < 0.01 or FDR ≤ 0.1 for significant enrichment.

For *Snhg26*-KD and -KO samples, transcriptome-level Pearson correlations were calculated using Microsoft Excel “CORREL” function. Pearson correlations were adjusted to show a gradient of differences between WT ESCs (set at 0) and NPCs (set to 1) for *Snhg26-*KD and between WT ESCs (set at 0) and EpiLCs (set to 1) for *Snhg26-*KO.

### Transcriptome assembly, isoform classification quantification, and alternative splicing analysis

Transcriptome assembly, isoform classification, gene and isoform quantification, and alternative splicing analysis was performed using TAMA (tc_version_date_2022_02_22)^117^, SQANTI 3 (v5.1.1)^118^, IsoQuant (v3.6.1)^121^, and SUPPA2 (v2.43) ^119^, respectively, as previously done in Fort *et al.* (in preparation).

### Data availability

Datasets from other referenced works can be accessed either in the NCBI Gene Expression Omnibus (GEO) repository using their GSE numbers or in the database of the EMBL’s European Bioinformatics Institute using their E-MTAB numbers (OKS reprogramming (Fort *et al.*, in preparation); RADICLseq^51^: GSE132192).

### Analysis RADICLseq data

Dataset^51^ from mouse ESCs cultured in DMEM supplemented with 15% FBS and LIF was downloaded as text files and analyzed on Microsoft Excel.

### *Snhg26* pulldowns

Chromatin Isolation by RNA Purification (ChIRP) was performed following Chu *et al.*, JOVE 2012. RNA pulldowns of *Snhg26* RNA interactors were adapted from Chu *et al.*, JOVE 2012 as follow: Wild-type and KO-ESC lines cultured in a humidified incubator at 37°C under 5% CO2 and maintained in 2i-LIF medium on 0.1% porcine gelatin (Sigma-Aldrich #G1890) were expanded to a total of 25 x 10^6^ cells. Cells were rinsed with DPBS (ThermoFisher Scientific #14190250) once and pre-incubated with the crosslinking solution (2:3 mixture of 2i-LIF medium and DPBS, supplemented with 25µg/mL 4’’-Aminomethyl-4,5’’,8-trimethylpsoralen (AMT, Cedarlane #17162-10)) for 30 minutes at 37°C. Cells were then dissociated using TrypLE^TM^ Express Enzyme (ThermoFisher Scientific #12483020) before counting and washing twice with DPBS. Next, cells were incubated in 2mL of DPBS supplemented with 25µg/mL AMT for 20 minutes on ice protected from light. Crosslinking was performed by exposing the cell suspension to 365nm UV (Stratalinker) in lidless plates for 30 minutes on ice, with shaking every 5 minutes. Cells were pelleted at 2000g for 5 minutes at 4°C and flash frozen. Total RNA was extracted using TRIzol^TM^ Reagent (ThermoFisher Scientific #15596018) according to the manufacturer’s instructions and quantified using Qubit^TM^ RNA broad range assay Kit (ThermoFisher Scientific #Q10210) on a Qubit ^TM^ 4 Fluorometer (ThermoFisher Scientific). For the pulldown, two sets of four different 3’ biotinylated DNA oligonucleotide probes targeting *Snhg26* transcript, or a control *GFP* transcript were designed manually and synthesized through IDT^TM^ or LGC Biosearch technologies (oligonucleotide probe sequences are described in **Table S4**). 100pmoles of pooled probes were hybridized to 120μg of total RNA in 4mL of Hybridization buffer (50mM Tris pH7.5 (Bio Basic #TB0194), 4mM EDTA, 1% SDS, 500mM NaCl, 10% formamide (Bio Basic #FDB0212), supplemented with 1mM PMSF (Bio Basic #PB0425), RNase inhibitor Murine 1% (New England Biolabs (NEB) #M0314L) and 1X Protease Inhibitor Cocktail (PIC, Sigma-Aldrich #11873580001)) for 16 hours at 37°C under agitation. 5μg of RNA was kept as input. RNA-probe complexes were then pulled-down by incubating 30 minutes at 37°C with 100μL of Hydrophilic magnetic streptavidin-coupled beads (NEB #S1421S) previously washed twice with 1mL of Lysis buffer (50mM Tris pH7.5, 10mM EDTA, 1% SDS, 1mM PMSF (Bio Basic #PB0425), RNase inhibitor Murine 1% (New England Biolabs #M0314L) and 1X Protease Inhibitor Cocktail (PIC, Sigma-Aldrich #11873580001)). Next, RNA-probe-bead complexes were washed 5 times 5 minutes at 37°C under agitation in a rotisserie incubator with 1mL of fresh Wash buffer (2X SSC buffer, 0.5% SDS and 1mM PMSF). RNA-probe complexes were eluted by incubating 45 minutes at 50°C under agitation in 100μL of PK buffer (10mM Tris pH7, 100mM NaCl, 1mM EDTA, 0.5% SDS, 1mg/mL proteinase K (Bio Basic #PB0451)). Eluates were heated 10 minutes at 95°C and retrieved from the beads after spinning 2 minutes at 16,000g. All along, beads were separated from solution using the DynaMag™-2 magnet (Fisher Scientific #12-321-D) and Eppendorf^TM^ LoBind microcentrifuge tubes (Fisher Scientific #13-698-791) were used to limit material loss. Finally, eluates were reversed-crosslinked by exposing them to 254nm UV (Stratalinker) for 15 minutes on ice, in lidless tubes, and RNA was extracted and concentrated using the RNA Clean & Concentrator^TM^-5 Kit (Zymo Research #76020-604) according to the manufacturer’s instructions. RNA inputs were quantified again using Qubit^TM^ RNA Broad Range Assay Kit. 25% of the *Snhg26* pulldown samples were analyzed by RT-qPCR to validate enrichment of *Snhg26*.

DNA pulldown samples were sequenced with long-read DNA sequencing using a Ligation Sequencing Kit (ONT #SQK-LSK109), with each sample barcoded with the Native Barcoding Expansion (ONT #EXP-NBD104), according to the manufacturer’s protocol: “Native barcoding amplicons (with EXP-NBD104, EXP-NBD114, and SQK-LSK109)” version NBA_9093_v109_revA_12Nov2019. Barcoded samples were pooled, loaded on a MinION sequencer (ONT) using one R9.4.1 Flow Cell (ONT #FLO-MIN106D), and sequenced using the MinKNOW^TM^ (v20.10.3) software.

RNA pulldown samples were DNase treated, reverse transcribed and PCR amplified for long-read RNA sequencing as explained above in the “Library preparation and long-read RNA-sequencing” section. Libraries were loaded on a MinION sequencer (ONT) using R9.4.1 Flow Cells (ONT #FLO-MIN106D) and sequenced using the MinKNOW^TM^ (v22.12.7) software, or on a PromethION 2 Solo (ONT) sequencer using R10.4.1 Flow Cells (ONT #FLO-PRO114M) and sequenced using the MinKNOW^TM^ (v23.07.15) software.

Reads were basecalled and oriented as explained in Fort *et al.* (in preparation) and mentioned in the “LrRNAseq data analysis of gene expression” section. Read CPM values from DNA pulldowns were obtained with the featureCounts tool^114^ from the Subread package (v2.0.6), and transcript TPM values from RNA pulldowns were obtained with the IsoQuant (v3.6.1) software^121^. For RNA pulldowns, *Snhg26* interactors specifically pulled-down with control *GFP* probes, *Snhg26-*specific even probes or *Snhg26-*specific odd probes in WT ESCs compared to KO-ESCs were established based on TPM fold change ≥ 1.5 (enriched in WT ESCs) or ≤ 0.67 (enriched in KO-ESCs) and p-value ≤ 0.05 (calculated by 2-tailed t-test with unequal variance). Genes of transcripts enriched with both *Snhg26-*specific probes and *GFP* control probes were filtered out.

GO-terms enrichment was assessed using the Database for Annotation, Visualization and Integrated Discovery (DAVID) (v2022q3)^116^. Biological Processes (BP) annotations were analyzed with a cut-off of p-value < 0.025 for significant enrichment.

### Analysis of repeat elements

We downloaded the RepeatMasker annotation file from the UCSC genome browser^122^. Read counts from *Snhg26* pulldowns were obtained with the featureCounts tool^114^ from the Subread package (v2.0.6). Enrichment over input was represented with SEM and p-values were calculated by two-tailed t-test (two samples with unequal variance) and a cut-off p-value < 0.05. Meta-repeat plots were generated with the deepTools software^123^ (v3.5.4), using the bamCompare command with default parameters to show enrichment of *Snhg26* pulldowns over control pulldowns, the computeMatrix command with the scale-regions argument and a bin size of 10, an d the plotHeatmap command to generate the metaplots.

### Expression of repeat elements whole transcriptome analysis

Read counts from *Snhg26-*KD ESCs were obtained with the featureCounts tool from the Subread package (v2.0.6)^114^. SINE elements were considered differentially expressed with fold-changes > 1.5 and p-values < 0.05 (calculated by two-tailed t-test, two samples with unequal variance). Differences between expected and observed numbers of up- and down-regulated SINE elements of a family were assessed with a Chi-squared test. n.s. ≥ 0.05, *** < 0.001, **** < 0.0001.

